# The Genetic Landscape of a Metabolic Interaction

**DOI:** 10.1101/2023.05.28.542639

**Authors:** Thuy N. Nguyen, Christine Ingle, Samuel Thompson, Kimberly A. Reynolds

## Abstract

Enzyme abundance, catalytic activity, and ultimately sequence are all shaped by the need of growing cells to maintain metabolic flux while minimizing accumulation of deleterious intermediates. While much prior work has explored the constraints on protein sequence and evolution induced by physical protein-protein interactions, the sequence-level constraints emerging from non-binding functional interactions in metabolism remain unclear. To quantify how variation in the activity of one enzyme constrains the biochemical parameters and sequence of another, we focused on dihydrofolate reductase (DHFR) and thymidylate synthase (TYMS), a pair of enzymes catalyzing consecutive reactions in folate metabolism. We used deep mutational scanning to quantify the growth rate effect of 2,696 DHFR single mutations in 3 TYMS backgrounds under conditions selected to emphasize biochemical epistasis. Our data are well-described by a relatively simple enzyme velocity to growth rate model that quantifies how metabolic context tunes enzyme mutational tolerance. Together our results reveal the structural distribution of epistasis in a metabolic enzyme and establish a foundation for the design of multi-enzyme systems.

## INTRODUCTION

Enzymes function within biochemical pathways; exchanging substrates and products to generate useful metabolites. This metabolic context places constraints on enzyme velocity — the product of both catalytic activity and enzyme abundance. For example, the relative velocities of some enzymes must be coordinated to avoid accumulation of deleterious metabolic intermediates^1–3^. In other instances, optimal enzyme abundance is set by a tradeoff between the cost of protein synthesis and the benefit of efficient nutrient utilization^4–6^. Considered at the pathway scale, metabolic enzymes are often produced in evolutionarily conserved stoichiometric ratios across species^7^, providing further indication that relative — not just absolute — enzyme velocity is under selection. More generally, the relationship between the velocity of a given enzyme, metabolic flux through a pathway, and cellular growth rate is non-linear and shaped by interactions amongst pathway enzymes (**Fig.1a**). Indeed, a key result of metabolic control theory is that the control coefficient of an enzyme — defined as the fractional change in pathway-level flux given a fractional change in enzyme velocity — depends on the starting (native) velocity of the enzyme, but *also* the velocity of all other enzymes in the pathway^8,9^. That is to say, given that enzymes act sequentially to produce metabolites, the effects of mutations on cellular phenotype can be buffered or amplified depending on which enzymatic reactions control metabolic flux. As a consequence, enzyme mutations that are neutral in one context may have profound consequences for metabolic flux and growth rate in the background of variation in another^10–13^. This context-dependence, or epistasis, amongst metabolic enzymes need not be mediated by direct physical binding, but emerges indirectly through shared metabolite pools and a need to maintain flux while avoiding the accumulation of deleterious intermediates^6,11,14^.

While much prior work has explored the constraints on protein sequence and evolution induced by physical protein-protein interactions, the sequence-level constraints emerging from these sorts of non-binding functional interactions in metabolism remain unclear. How is this biochemically-mediated epistasis organized in the protein structure and reflected in the sequence? A quantitative understanding of how pathway context shapes sequence and activity would assist in the interpretation of disease-associated mutations, the design of new enzymes, and directing the laboratory evolution of metabolic pathways. To begin to address this, we examined the residue-level epistatic interactions between a pair of enzymes that catalyze consecutive reactions in folate metabolism: dihdyrofolate reductase (DHFR) and thymidylate synthase (TYMS). The activity of these enzymes is strongly linked to *E. coli* growth rate, they are frequent targets of antibiotics and chemotherapeutics, and our prior work showed that they co-evolve as a module both in the laboratory and across thousands of bacterial genomes^1^. Taking this enzyme pair as a simplified model system in which to examine a biochemically-mediated epistatic interaction, we created a mathematical model relating variation in DHFR and TYMS catalytic parameters to growth rate using a focused set of well-characterized point mutants. Then, to more deeply test this model and comprehensively map the pattern of epistasis between these two enzymes, we measured the effect of nearly all possible DHFR single mutations (2,696 in total) in the context of three TYMS variants selected to span a range of catalytic activities. The model predicted – and the data showed – that TYMS background profoundly changed both the sign (buffering vs. aggravating) and magnitude of DHFR epistasis. Mapping the epistatic effects of mutation to the DHFR tertiary structure revealed that they are organized into distinct clusters based on epistatic sign. Additionally, mutations with the largest magnitude epistatic effect to TYMS centered around the DHFR active site, while more weakly epistatic positions radiated outwards. Finally, we inferred approximate values for DHFR catalytic power (*k_cat_*/K_m_) across all 2,696 mutations by using the growth rate measurements across TYMS backgrounds to constrain the enzyme velocity to growth rate model. The residues linked to catalysis form a structurally distributed network inside the enzyme and are highly evolutionarily conserved. Taken together, our data demonstrates at single-residue resolution how epistasis mediated through a biochemical interaction reshapes a mutational landscape. Our results indicate that metabolic context can strongly modulate enzyme evolution in both the clinic and the lab by facilitating or frustrating available mutational paths. More generally, our results invite one to consider new ideas for the joint design of multi-enzyme systems that take into account shared constraints on relative activity and sequence.

## RESULTS

### An enzyme velocity to growth rate model for DHFR and TYMS

We constructed a mathematical model relating changes in DHFR and TYMS catalytic parameters to growth rate phenotype. Our goals were to (1) formalize our previous empirical findings describing DHFR/TYMS biochemical coupling^1^, (2) quantify the absolute and relative constraints on DHFR and TYMS catalytic activity, and (3) define the relationship between biochemical activity and epistasis. DHFR and TYMS play a central role in folate metabolism, a well-conserved biochemical pathway involved in the synthesis of purine nucleotides, thymidine, glycine, and methionine^15^ (**Fig. 1b**). Consequently, this pathway is strongly linked to cell growth and a frequent target of antibiotics and chemotherapeutics. DHFR is a 159 amino acid enzyme that catalyzes the reduction of dihydrofolate (DHF) to tetrahydrofolate (THF) using NADPH as a cofactor. The reduced THF then serves as a carrier for activated one-carbon units in downstream metabolic processes. TYMS catalyzes the oxidation of THF back to DHF during deoxythymidine synthesis and is the sole enzyme responsible for recycling the DHF pool^16,17^. Prior work from ourselves and others indicates that these two enzymes are strongly functionally coupled to each other and less coupled to the remainder of the pathway: they co-evolve in terms of synteny and gene presence-absence across bacterial species^1^, inhibition of DHFR with trimethoprim is rescued by suppressor mutations in TYMS in both the lab and the clinic^1,18^, and loss-of-function mutations in DHFR are rescued by loss-of-function mutations in TYMS^1,19,20^. Metabolomics data indicated that loss of DHFR function resulted in accumulation of DHF and depletion of reduced folates, and that compensatory loss of function mutations in TYMS help to restore DHF and THF pools to more native-like levels^1,21,22^. Thus, DHFR and TYMS are a growth-linked two-enzyme system where epistasis is driven by a biochemical interaction, with the added simplification that they are relatively decoupled from surrounding metabolic context.

**Figure 1.**
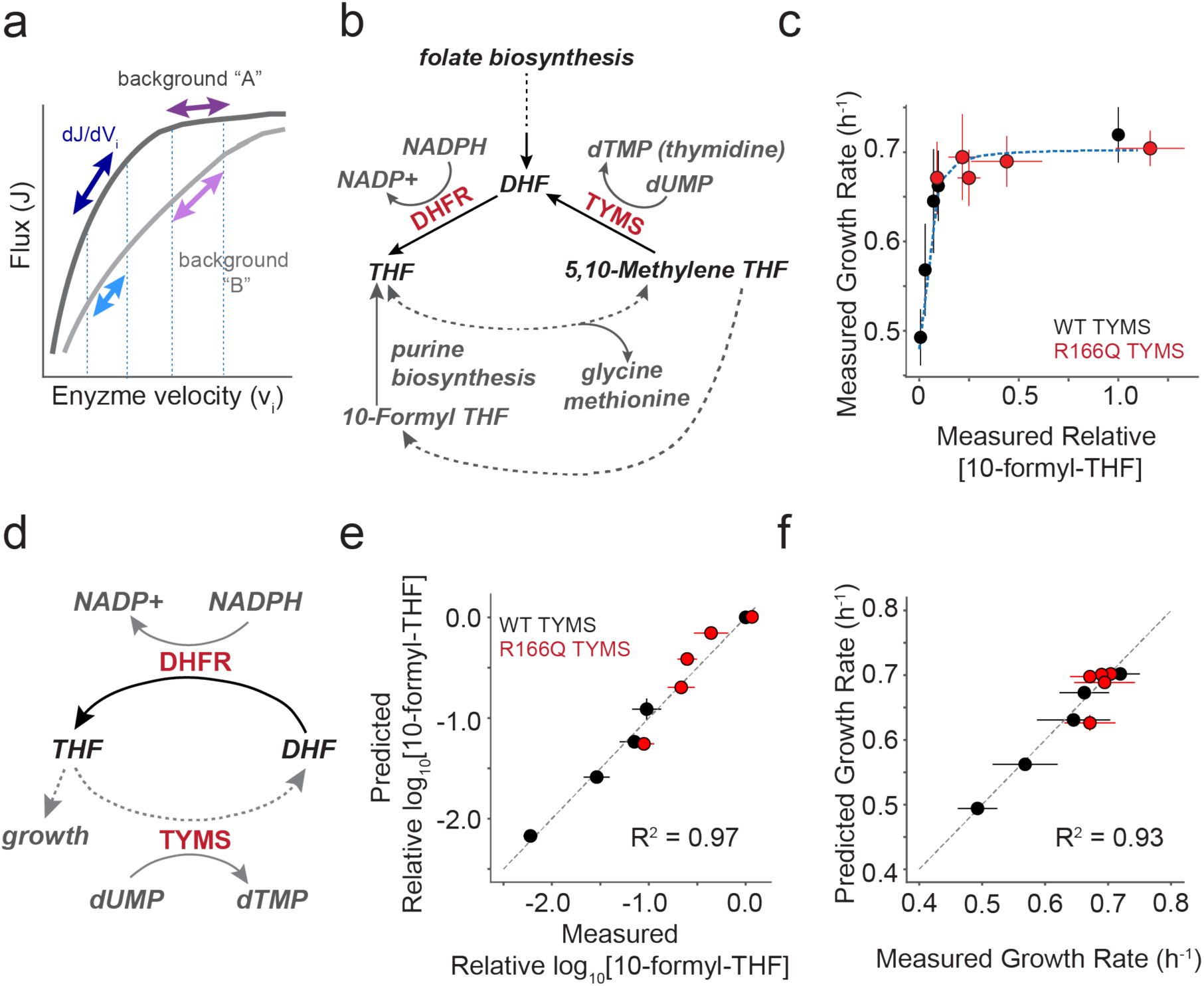
Constructing a biochemistry-to-growth model for DHFR and TYMS. **a)** Schematic describing the relationship between metabolic pathway flux and enzyme velocity. Many enzymes show a hyperbolic relationship between velocity and flux; the enzyme control coefficient describes the fractional change in flux given a fractional change in velocity. Control coefficients vary with the starting enzyme velocity (purple and dark blue arrows, background “A”) and can change with genetic background (violet and light blue arrows, background “B”). Consequently mutations can have a strong effect on flux and growth in one context but not in another (compare violet and purple arrows). **b)** The role of DHFR and TYMS in folate metabolism. Metabolites are labeled in grey or black italic text. Dotted lines indicate multiple intermediate reactions that are summarized with a single line. **c)** The relationship between the experimentally measured relative abundance of [10-formyl-THF] and *E. coli* growth rate. Red points indicate five DHFR variants in the background of TYMS R166Q (a near catalytically inactive variant) and black indicates the same DHFR variants in the context of WT TYMS. Error bars indicate the standard deviation across N=3 replicates for both growth rate (y-axis) and 10-formyl-THF abundance (x-axis). The blue dotted line indicates the best fit for a hyperbolic model (Equation 1) relating THF abundance to growth. **d)** A simplified, abstracted version of the DHFR and TYMS system. Again dotted lines indicate multiple intermediate reactions that are summarized with a single line. **e)** The correlation between experimentally measured log_10_[10-formyl-THF] relative abundance and the model prediction (as computed with Equation 3). The grey dotted line indicates x=y. Color coding is identical to **c**. Error bars in x indicate the standard deviation across N=3 replicate experimental measurements, error bars in y describe the standard deviation across ten fits obtained by jackknife (leave-one-out) sub-sampling the data and refitting the model. **f)** The correlation between experimentally measured and predicted growth rates for five DHFR point mutations in two different TYMS backgrounds (same mutants as in **c**,**e**). The grey dotted line indicates x=y. Color coding is identical to **c**. Error bars in x indicate the standard deviation across N=3 replicate experimental measurements, error bars in y describe the standard deviation across ten fits obtained by jackknife (leave-one-out) sub-sampling the data and refitting the model.

We first developed our mathematical model using a previously collected set of metabolomics and growth rate data for five DHFR point mutants in the background of both WT TYMS and TYMS R166Q (**Table S1, Table S2**). TYMS R166Q is an active site mutation with near complete loss of catalytic activity^23^. The model comprises two parts: the relationship between intracellular THF abundance and growth rate, and the relationship between enzymatic activity and intracellular THF. First, we considered the relationship between intracellular THF abundance and growth rate as measured across all ten DHFR/TYMS sequence combinations. THF limitation restricts the production of several growth-linked factors, including thymidine, methionine, glycine, and the purine precursors inosine and AICAR. Under the experimental conditions of our growth rate assays — M9 minimal media with 0.4% glucose, 0.2% amicase, and 50 µg/ml thymidine — thymidine is not growth limiting (TYMS R166Q is rescued to WT-like growth) and amicase provides a source of free amino acids. These conditions — which remove selection pressure on TYMS due to thymidine production — emphasize coupling between DHFR and TYMS through the shared THF pool which must be used to produce purine nucleotides. We previously observed a hyperbolic dependence of growth rate on reduced folate abundance for many THF species in these experimental conditions^1^. We selected 10-formyl THF with three glutamates as a representative growth-linked reduced folate for modeling given it’s clear relationship to growth and proximity to purine biosynthesis. Following a similar approach as Rodrigues et al, we fit a single four-parameter sigmoidal function relating growth rate to the experimental measurements of intracellular THF concentration^24^ (**Fig. 1c**).

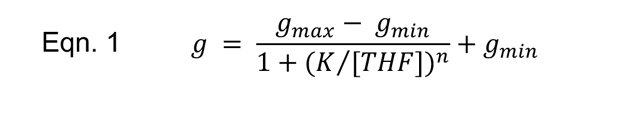

Here, *g*_*max*_ represents the maximal growth rate, *g*_*max*_ is the minimal growth rate, *K* is a constant that captures the concentration of THF that yields 50% growth, and *n* is a Hill coefficient (**Table S3**).

Next, we sought to connect variation in DHFR and TYMS enzyme velocity to intracellular THF concentrations. To simplify our model, we reduced the pathway to a cycle in which DHFR and TYMS catalyze opposing oxidation and reduction reactions (**Fig. 1d**). This abstraction assumes that DHFR and TYMS dominate turnover of the DHF and THF pools, and that the reduced folates are considered as a single THF pool. While this simplification clearly omits much of folate metabolism, it allows us to write a rate equation that isolates the recycling of THF in terms of a small number of measurable biochemical parameters:

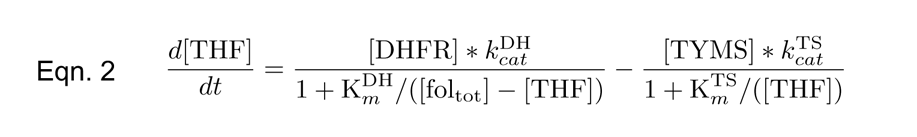

In this equation, DHFR and TYMS are treated as catalyzing opposing reactions with Michaelis Menten kinetics, providing a relationship between steady state kinetics parameters (*k*^DH^_*cat*_, *k*^DH^_*m*_, *k*^TS^_*cat*_, *k*^TS^_*m*_) and intracellular THF abundance. From this equation one can find an analytical solution for the steady state concentration of THF in the form of the Goldbeter-Koshland equation ^25,26^.

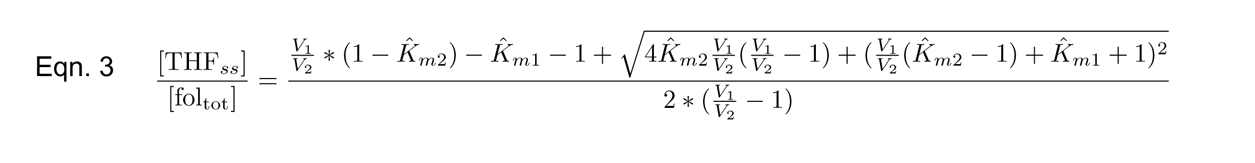

Where:

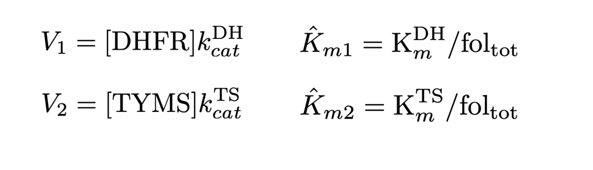

In our initial model construction, the steady state catalytic parameters (*k*^DH^_*cat*_, *k*^DH^_*m*_, *k*^TS^_*cat*_, *k*^TS^_*m*_) were experimentally measured *in vitro* using purified samples of all mutants, with the exception of TYMS R166Q which is near-inactive and assigned an arbitrarily low *k_cat_* and high K_m_ (**Table S1,S2**). Four fit parameters remain in Equation 3: (1) the concentration of the total folate pool ([fol_-.-_]) (2) the intracellular concentration of DHFR ([DHFR]), which we treated as identical across all variants (as our model will eventually describe thousands of DHFR mutations, and we wished to avoid overparameterization), (3) the intracellular concentration of WT TYMS ([TYMS_/+_]), and (4) the intracellular concentration of TYMS R166Q (>TYMS_01223_?). This relatively simplified model showed good correspondence to the data when fit (R^2^ = 0.96, **Fig. 1e, Table S3**). Equations 1 and 3 were then combined to estimate growth rate as a function of both DHFR and TYMS activity, by linking catalytic activity to THF abundance, and then THF abundance to growth rate. The complete model worked well to predict growth rate on our initial training set (**Fig. 1f**).

### The growth rate effect of DHFR mutations changes magnitude and sign depending upon TYMS background

To more rigorously test our model and understand its’ predictions, we expanded our dataset to include more DHFR and TYMS variants with experimentally characterized activities. As our initial model was developed using only two extreme TYMS variants (wild-type and a near complete loss of function variant, R166Q), we were particularly curious to evaluate model performance for TYMS mutations with intermediate effects on catalysis and *E. coli* growth. We identified candidate TYMS mutations by examining an earlier growth complementation study^27^. A handful of these mutants were then cloned, screened for expression, and when possible, purified and characterized. Through this mini-screen we selected two mutations that stably expressed, purified robustly, and yielded intermediate activities: TYMS R127A and Q33S (**Fig. 2a**). The R127A mutation is located in the TYMS active site and is one of four arginines that coordinate the substrate (dUMP) phosphate group. The Q33S mutation is located at the TYMS dimer interface, distal to the active site. We observed that R127A was more deleterious to catalytic function than Q33S, but that both mutations were more active than R166Q (which shows almost no measurable activity *in vitro*, **Fig. 2b, Table S2**).

**Figure 2.**
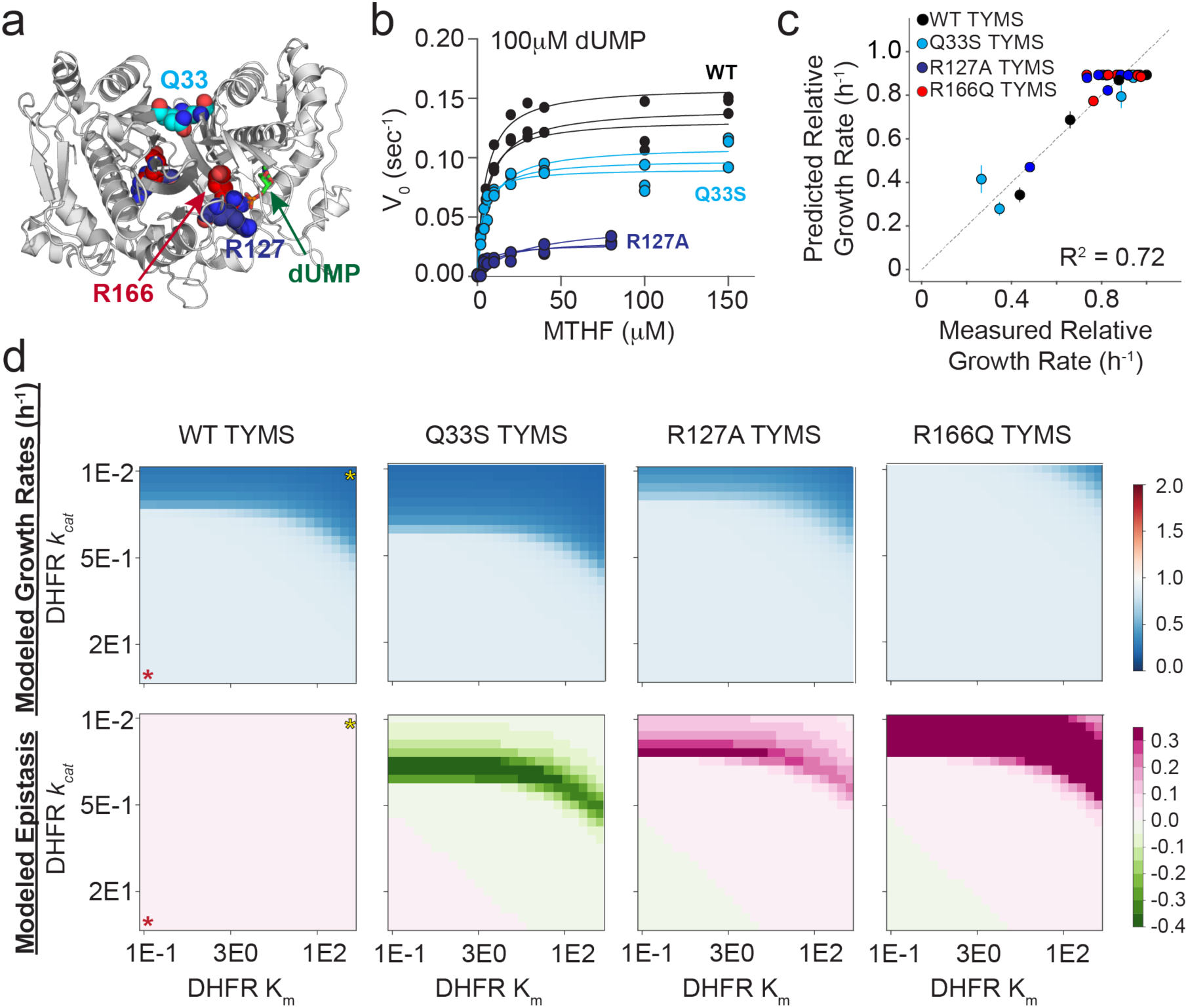
TYMS catalytic activity modulates the epistatic constraints on DHFR sequence and catalytic activity. **a)** Location of the TYMS point mutations (PDBID: 1BID^60^). TYMS functions as an obligate domain-swapped homodimer; active sites include residues from both monomers (white and grey cartoon). Positions mutated in this study are in colored spheres, and indicated with arrows (Q33 – cyan, R127 – navy, R166 – red). The dUMP substrate is in sticks, labeled and colored green. **b)** Michaelis Menten enzyme kinetics for WT TYMS (black), TYMS Q33S (cyan), and TYMS R127A (navy). Experimental replicates (3 total) are plotted individually. Points indicate experimental data and lines the best fit steady state model. **c)** Correlation between experimentally measured and model-predicted relative growth rates for seven DHFR variants in four TYMS backgrounds. Each point represents one DHFR/TYMS combination. Error bars in the x direction are SEM across triplicate growth rate measurements, error bars in y are the SEM estimated from jackknife (leave-one-out) sub-sampling the data and refitting the model. **d)** Heatmaps of simulated growth rates (top row) and epistasis (bottom row) as computed over a range of DHFR kinetic parameters in four TYMS backgrounds. In the left-most column of heatmaps a red star marks the highest activity enzyme (low K_m_, high *k_cat_*), while a yellow star marks the lowest activity enzyme. Growth rates are indicated with a blue-white-red color map, where a relative growth rate of one (white) is equivalent to WT. Epistasis values are indicated with a green-white-pink color map, where zero epistasis is shown in white.

We measured growth rates for seven catalytically characterized DHFR variants (a set of single and double mutants selected to span a range of catalytic activities) in the background of these four TYMS mutants (WT, R127A, Q33S and R166Q). The point mutants were created in a plasmid encoding both DHFR and TYMS (see Methods for details), and the plasmids were transformed into an E. coli selection strain lacking the endogenous DHFR and TYMS genes (ER2566 *ΔfolA ΔthyA*). Growth rates were measured in triplicate using a plate-reader-based assay (28 measurements total; **Fig. 2c**, **Fig S1a,c**). We used this focused dataset to re-parameterize the model equations, this time fitting five total parameters ([fol_tot_], [DHFR], [TYMS_WT_], >TYMS_WT_, [TYMS_R127A_], [TYMS_R166Q_], **Table S3**). This second round of fitting tested the ability of growth rate data alone to constrain the model — an important step because metabolomics data are available for only a limited number of DHFR and TYMS mutants and are inherently far lower throughput to collect than growth rates. This iteration of parameterization also tested the capacity of the model to capture TYMS mutations with intermediate effects on activity. The data were again well described by the model (**Fig. 2C, Fig. S1b,d**). We observed some variation in the fit parameters (relative to the older data in Fig.1); this difference might be attributed to the fact that our newer experiments used a revised selection vector backbone. As a control for overfitting, we tested the ability of the model to predict growth rates for arbitrary catalytic data. We randomly shuffled the catalytic parameters (*k_cat_* and K_m_) among mutations for both DHFR and TYMS, refit all free model parameters, and calculated the RMSD and R^2^ values between the best fit model and the shuffled data. Importantly, the model was generally unable to describe the experimental growth rate data when catalytic parameters were shuffled across both DHFR and TYMS (**Fig S1e,f**). This indicated that the model provided a specific description of our experiment and was not trivially overfit. The model was less sensitive to shuffling TYMS catalytic parameters (presumably because we included fit parameters describing the abundance of each TYMS mutation that can compensate for this shuffling, **Fig. S1h**). However, it was strongly sensitive to shuffling DHFR parameters (**Fig. S1g**). Taken together, this analysis indicated that the model provides a good description of the enzyme-velocity-to-growth-rate relationship and can be used to predict and interpret how molecular changes in DHFR and TYMS activity modulate growth rate phenotype.

To examine the model more closely, we considered the relationship between TYMS catalytic activity and DHFR mutational sensitivity. As in previous work, we observed that loss-of-function mutations in DHFR can be partly or even entirely rescued by the loss-of-function mutation TYMS R166Q in the presence of thymidine (**Fig. S1a,c**). TYMS R127A, a less severe loss of function mutation, showed a similar albeit more modest trend – this mutation was able to partly rescue growth for some (though not all) DHFR mutations. A central factor behind DHFR and TYMS biochemical coupling is that loss-of-function mutations in TYMS help to preserve reduced folate pools, allowing THF to shuttle one-carbon units in downstream biochemical processes like purine biosynthesis even when DHFR activity is low. Moreover, loss of TYMS activity reduces accumulation of DHF, which can inhibit upstream reactions^1,22^. Thus, the TYMS R166Q and R127A variants show positive (buffering) epistasis to low-activity DHFR mutations. In contrast to our expectation that a more intermediate mutation would also demonstrate intermediate levels of buffering epistasis, TYMS Q33S shows negative (or amplifying) epistasis to some DHFR mutations (**Fig. S1c**). This means that these DHFR mutations are more deleterious in the background of TYMS Q33S than in the native TYMS context. Our model accounted for this observation by increasing the intracellular concentration of TYMS Q33S by three fold (a fit parameter, **Table S3**) such that the V_max_ of TYMS Q33S becomes greater than wildtype ([TYMS_Q33S_]*k*_*cat*_^TYMS_Q33S^ > [TYMS^WT^]*k*^TYMS_WT^_*cat*_). This in turn increased the intracellular requirement for DHFR activity, resulting in negative epistasis.

To further explore the pattern of epistasis across TYMS backgrounds, we simulated growth rates over a range of DHFR *k_cat_* and K_m_ values in each TYMS background (**Fig. 2d**). This provided a comprehensive prediction of the TYMS-induced constraints on DHFR activity. In particular, we obtained a regime of DHFR *k_cat_* and K_m_ values that is sufficient to support growth for each TYMS mutation. From these data we computed epistasis. These results indicated that TYMS Q33S has negative epistasis to DHFR variants spanning a well-defined band of catalytic parameters. The simulations also indicated that R127A has weak positive epistasis over a regime of moderately impaired DHFR variants, but is insufficient to rescue growth for the strongest loss of function variants. Finally, TYMS R166Q was observed to be broadly rescuing; DHFR variants need only a negligible amount of activity to support growth in this context. Together, our simulations showed that the sign and magnitude of DHFR epistasis are strongly tuned by TYMS background, and provided quantitative predictions of the catalytic regimes where epistasis is most apparent.

### The single-mutant landscape of DHFR is strongly modulated by TYMS context

Next we examined the structural pattern of biochemical epistasis at the residue level across DHFR. This also provided an opportunity to see if the model predictions — negative epistasis for Q33S and broadly positive epistasis for R166Q — held true across a larger dataset. To accomplish this, we created a plasmid-based saturation mutagenesis library of DHFR containing all possible single mutations at every position (3002 total). This library was subcloned into all three TYMS backgrounds. Sequencing showed that these libraries are well-distributed and approach full coverage of all single mutations (97.1% - WT TYMS, 94.6% - TYMS Q33S, 99.3% -TYMS R166Q) (**Fig. S2**). We transformed these libraries into the *E. coli* selection strain (ER2566 *ΔfolA ΔthyA*). Transformants for each library were then grown as a mixed population in selective media in a turbidostat to ensure maintenance of exponential growth and constancy of media conditions. By quantifying the change in the relative frequency of individual mutants over time with next generation sequencing, we obtained a growth rate difference relative to WT DHFR for nearly all mutations in the library (**Fig. 3a**, **Table S4**, see methods for details). All relative growth rate measurements were made in triplicate, with good concordance among replicates (**Fig. S3**).

**Figure 3.**
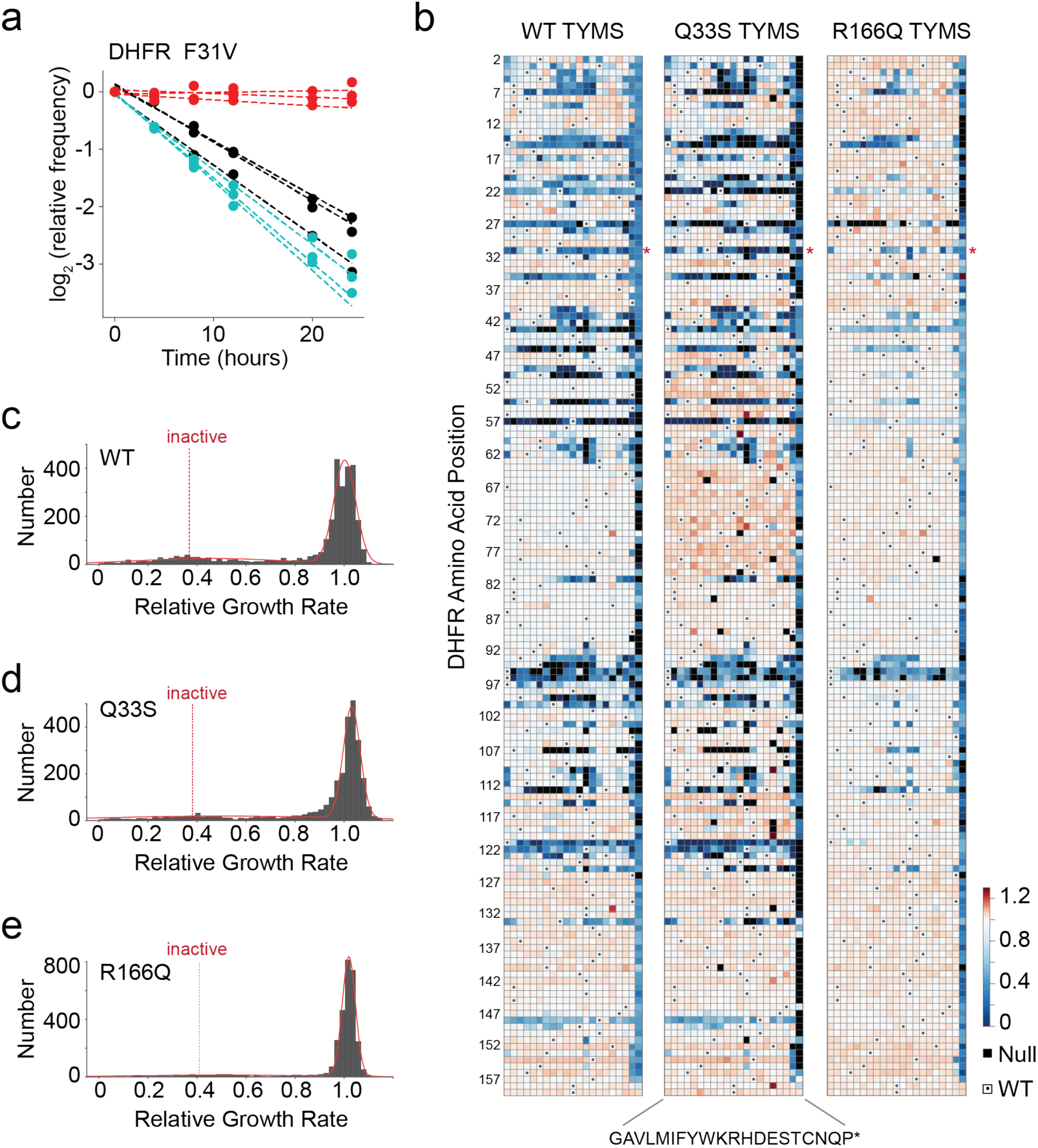
The effects of DHFR mutation on growth rate in three TYMS backgrounds. **a)** Sequencing-based growth rate measurements for DHFR F31V in three TYMS backgrounds: R166Q (red), Q33S (cyan), and WT (black). Each point represents one triplicate experimental measurement. Dotted lines indicate linear regression fits to each replicate, the slope of each line is the inferred growth rate (relative to WT) for that DHFR/TYMS mutant combination. **b)** Heatmaps of the growth rate effect for all DHFR single mutations. DHFR positions are along the horizontal axis; amino acid residues (along the vertical axis) are organized by physiochemical similarity. The displayed relative growth rate is an average across three replicates, and is normalized such that the WT DHFR is equal to one. Red indicates mutations that increase growth rate, white indicates mutations with wild-type like growth, and blue indicates mutations that decrease growth rate. Null mutations (black squares) were not observed by sequencing after the first two time points, and thus there was insufficient data for growth rate inference. Small dots mark the WT residue identity in each column. **c)** The distribution of DHFR mutational effects in the WT TYMS background. The red line indicates a best-fit double gaussian, grey bars are the data. The red, dashed “inactive” line marks the average relative growth rate for nonsense mutations (stop codons) in the first 120 positions of DHFR. The WT DHFR growth rate is equal to one. **d)** The distribution of DHFR mutational effects in the TYMS Q33S background, color coding identical to (**c**) **e)** The distribution of DHFR mutational effects in the TYMS R166Q background, color coding identical to (**c**). Note that the y-axis for (**e**) is distinct from (**c**) and (**d**).

The entire dataset showed that the DHFR mutational landscape was strongly dependent on TYMS background (**Fig. 3b-e**). In all three TYMS backgrounds, the distribution of growth rate effects was bi-modal and reasonably well-described by a double gaussian containing one peak of near-neutral mutations and another (far smaller) peak of mutations with highly deleterious growth rate effects. This is the expected result for an enzyme that shows a sigmoidal relationship between activity and growth. In the native TYMS context, the vast majority of mutations fall into the near-neutral peak. However, there is a substantial fraction (12%, 343 total) that display growth rates at or below that of “inactive”, where “inactive” was defined as the average growth rate across nonsense mutations in the first 120 residues of DHFR. Consistent with expectation, mutations at known positions of functional importance tended to be deleterious in the WT TYMS context (W22, D27, F31, T35, L54, R57, T113, G121, and D122)^28^. For example, both W22 and D27 are directly in the active site and serve to coordinate substrate through a hydrogen bonding network^29^, G121 and D122 are part of the βf-βg loop and stabilize conformational changes associated to catalysis^30,31^, and F31 contacts the substrate and is associated to the “network of promoting motions”^32,33^. In the TYMS Q33S context, many of these deleterious mutations had even more severe effects or were classified as “Null”. Null mutations disappeared from our sequencing counts within the first three time points (8 hours) of the selection experiment, preventing accurate inference of growth rate. For example, mutations at position 22 are deleterious in the WT TYMS context, and appear as Null or very deleterious in the Q33S context. The same pattern can be readily observed for positions 7,14,15, 22, 27, 31, 35, and 121. Again we saw that 12% of mutations have growth rates at or below that of “inactive” variants. Finally, in the TYMS R166Q context, there are very few deleterious mutations. Nearly all mutations are contained in the near-neutral peak, including mutations at highly conserved active site positions like M20, W22, and L28. Stop codons and mutations at the active site residue D27 continued to be deleterious, indicating that DHFR activity was still under (very weak) selection in the TYMS R166Q background. Nonetheless, only 5% of mutations displayed growth rates at or below those of inactive mutations. Thus TYMS R166Q is broadly buffering to DHFR variation.

To quantify the context dependence of mutational effects, we computed epistasis for all DHFR mutations with measurable relative growth rates in each of the three TYMS backgrounds (2,696 in total, see also methods) (**Fig. 4, Fig. S4, Table S5**). We assessed the statistical significance of epistasis by unequal variance t-test under the null hypothesis that the mutations have equal mean growth rates in both TYMS backgrounds. These p-values were compared to a multiple-hypothesis testing adjusted p-value determined by Sequential Goodness of Fit (P = 0.035 for TYMS Q33S and P = 0.029 for TYMS R166Q, **Fig. 4a,b**) ^34^. In the TYMS Q33S background, 95 mutations (3%) showed significant negative epistasis and 280 mutations (9%) showed significant positive epistasis. Many of the DHFR mutations with positive epistasis to Q33S were near-neutral in the WT context, and displayed small improvements in growth rate that were highly significant due to the low experimental error for the best-growing mutations (**Fig. 4c**). In contrast, the mutations with negative epistasis exhibited a range of growth rate effects in the WT context. For the TYMS R166Q background the overall proportion of significant epistatic mutations was larger: while only 41 mutations (1%) showed significant negative epistasis, 851 mutations (28%) showed significant positive epistasis. A smaller number of deleterious mutations in the WT context were not rescued by R166Q (grey points with relative growth rates near 0.25). One possibility is that these mutations are deleterious for reasons beyond disrupting metabolic flux; for example they may result in protein aggregation or off target physical interactions. Regardless, direct comparison of the relative growth rates of mutations across the WT, Q33S, and R166Q TYMS backgrounds makes it very obvious that TYMS R166Q was broadly rescuing, while TYMS Q33S had a more subtle effect that sometimes yielded negative epistasis (**Fig. 4c,d**).

**Figure 4.**
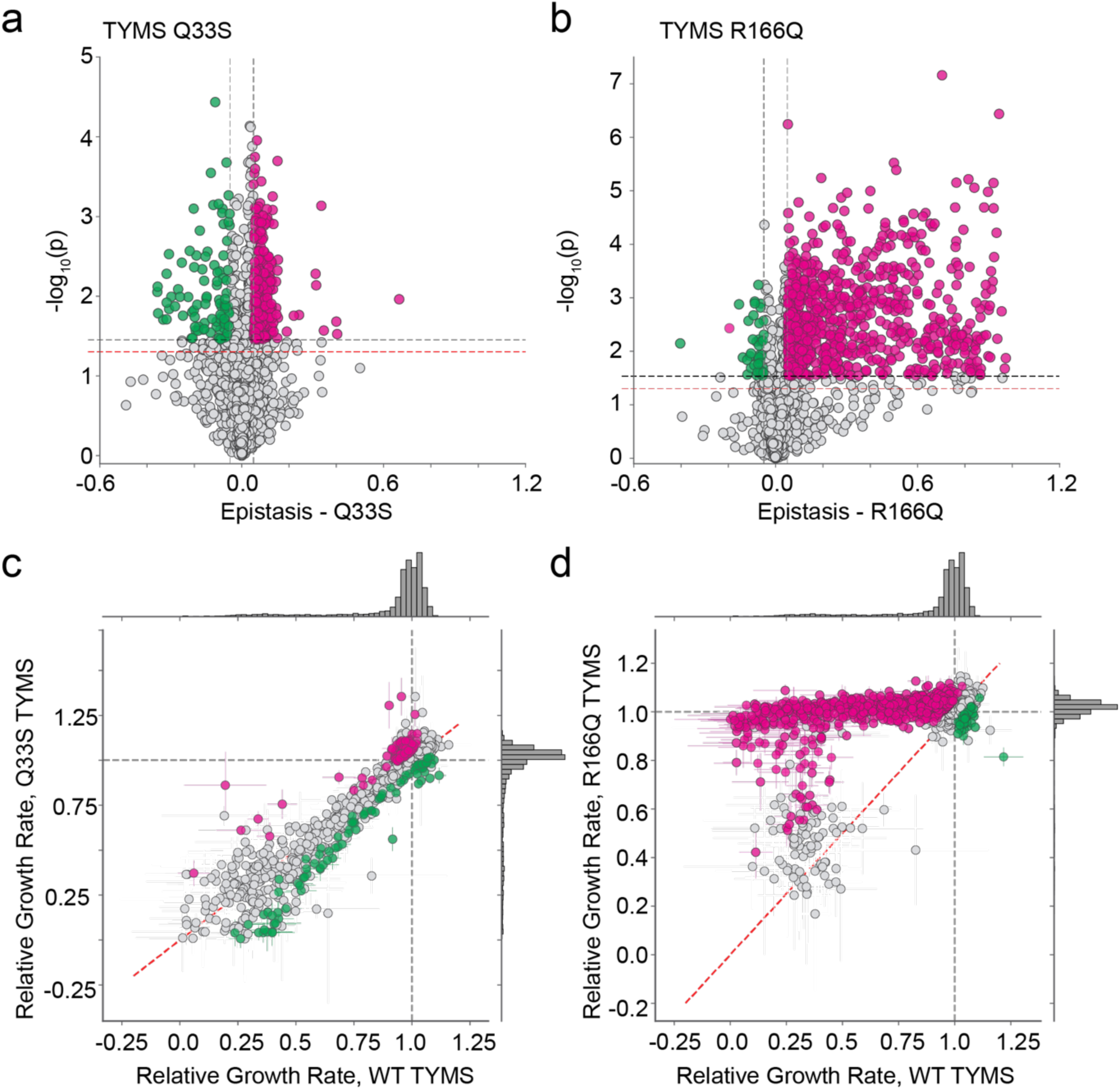
Epistatic coupling of DHFR to two TYMS backgrounds. **a)** Volcano plot examining the statistical significance of epistasis across all DHFR point mutations in the Q33S background. P-values were calculated by unequal variance t-test under the null hypothesis that the mutations have equal mean growth rates in both TYMS backgrounds (across triplicate measurements). The red horizontal dashed line marks the standard significance cutoff of P=0.05, the black horizontal dashed line indicates a multiple-hypothesis testing adjusted p-value (P=0.035). The grey vertical dashed lines indicate an empirical threshold for epistasis. Pink and green indicate statistically significant positive and negative epistasis respectively. **b)** Volcano plot examining the statistical significance of epistasis across all DHFR point mutations in the R166Q background. P-values were calculated as in (**a**); the multiple-hypothesis testing adjusted p-value for the R166Q background was (P=0.029). Color coding is identical to (**a**). **c)** Comparison of the relative growth rate effects for DHFR single mutants in the WT and TYMS Q33S backgrounds. The marginal distribution of growth rate effects is shown along each axis. Mutations with statistically significant positive and negative epistasis are indicated in pink and green respectively. The WT relative growth rate equals one, and is indicated with a dashed grey line across each axis. The dashed red line marks x=y. Error bars (in x and y) indicate standard deviation across three experimental measurements. **d)** Comparison of the relative growth rate effects for DHFR single mutants in the WT and TYMS R166Q backgrounds. Plot layout and color coding is identical to (**c**). Error bars (in x and y) indicate standard deviation across three experimental measurements.

### The enzyme velocity to growth-rate model captures the observed fitness landscapes and allows global estimation of mutational effects on catalysis

Next we sought to further test our enzyme velocity to growth-rate model using the deep mutational scanning data. We refit the model a third time, in order to include all available experimental information in parameterizing the model for this larger set of mutants. This included a larger dataset of 34 DHFR single mutants with previously reported *k_cat_* and K_m_ values. We additionally characterized *k_cat_* and K_m_ for four new DHFR mutations (I5K, V13H, E17V and M20Q) that exhibited strong sign epistasis to TYMS to more completely test our ability to predict epistasis. Together this yielded a set of 114 growth rate measurements with matched *k_cat_* and K_m_ values for DHFR and TYMS (38 DHFR mutations in 3 TYMS backgrounds, **Table S1**). We used these data to perform a bootstrap analysis; iteratively subsampling the data and refitting the model 1000 times to obtain standard deviations in our model fit and the eight associated parameters (**Fig. 5a**). The inferred parameters for this large set of sequencing-based growth rate measurements were qualitatively similar to those obtained for the smaller set of 28 plate-reader based growth rate measurements (7 DHFR mutants in 4 TYMS backgrounds, **Fig. 2**), but we observed some discrepancy in the estimated total folate pool and intracellular concentrations of TYMS (**Table S3**). In general, the total folate pool and concentrations of TYMS R166Q were more variable across the bootstrap analysis, indicating that these two parameters are less well constrained by our data (as indicated by the estimated variances in **Table S3**). Nevertheless, the inferred parameters in both the plate reader and sequencing-based experiments suggested similar relative expression levels of DHFR and the three TYMS point mutants, with Q33S being roughly 2-3 times more abundant than WT DHFR. Overall both the predicted growth rates and pattern of epistasis showed good agreement to our experimental observations (**Fig. 5a,b**).

**Figure 5.**
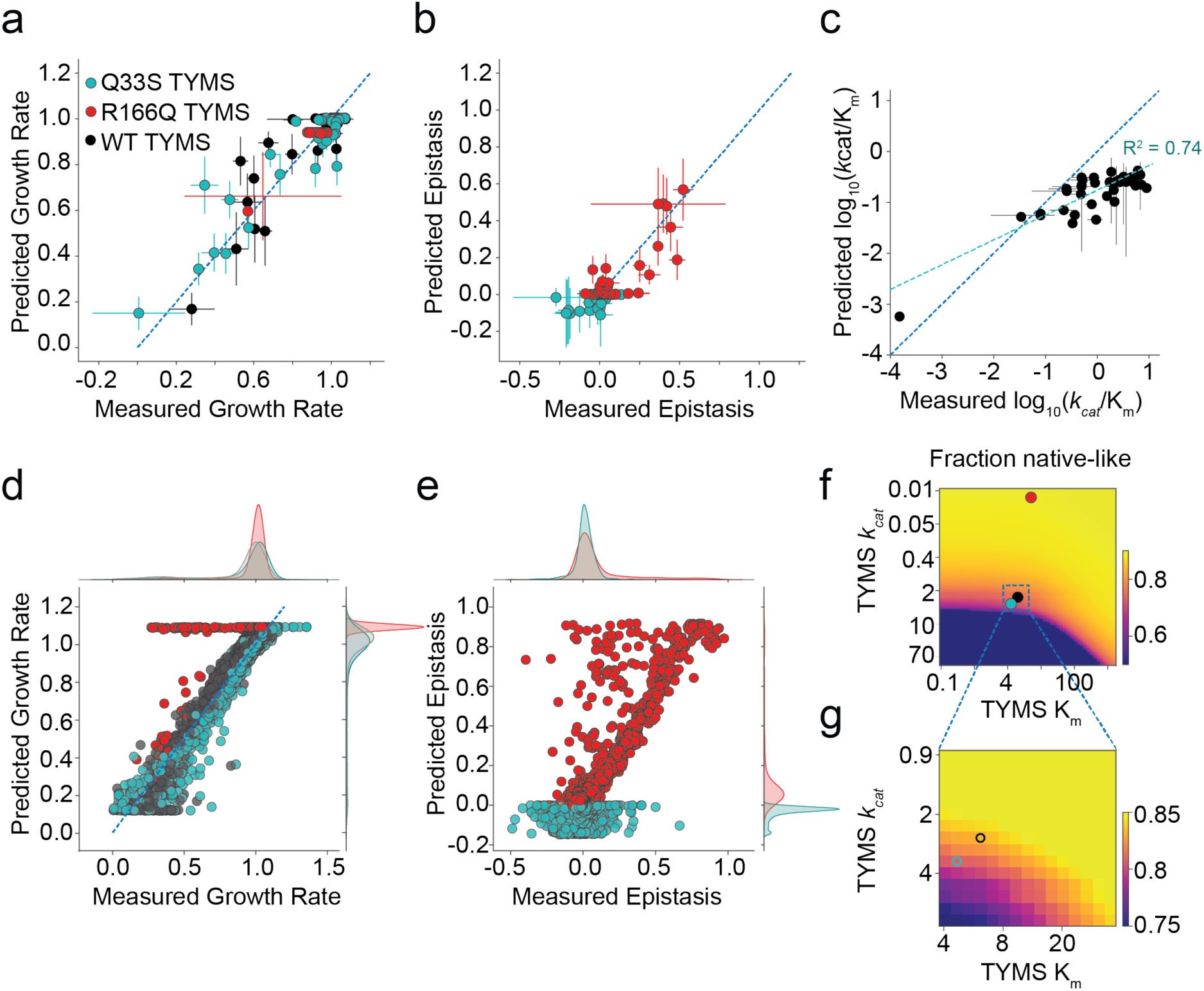
Global comparison of the biochemistry-to-growth model and deep mutational scanning data set. **a)** Correlation between the experimentally measured and predicted growth rates of 114 DHFR/TYMS mutant combinations (circles colored according to TYMS background). Horizontal error bars indicate standard deviation in experimentally measured growth rates across three replicate measurements, vertical error bars are the standard deviation in the predicted growth rates estimated by performing 1000 bootstrap re-samplings (and model fits) of the data. The dashed grey line indicates y=x. **b)** Correlation between the experimentally measured and model-predicted epistasis, as computed from the growth rate data in (**a**). Again, color coding indicates TYMS background (identical to **a**). The dashed grey line indicates y=x. Horizontal error bars indicate standard deviation propagated from the experimentally measured growth rates across three replicate measurements, vertical error bars are the standard deviation in the predicted epistasis values estimated by performing 1000 bootstrap re-samplings (and model fits) of the data. **c)** Correlation between the experimentally measured and computationally inferred log_10_(*k_cat_*/K_m_) values for 38 mutants of DHFR. Horizontal error bars describe the standard deviation across triplicate experimental measurements, vertical error bars indicate the standard deviation across 50 iterations of stochastic (Monte-Carlo based) model inference. **d)** Correlation between experimentally measured and predicted growth rates across the entire deep mutational scanning dataset. The marginal distribution of growth rate effects is shown along each axis. **e)** Correlation between experimentally measured and predicted epistasis across the entire deep mutational scanning dataset. The marginal distribution of epistatic effects is shown along each axis. **f)** Mutational tolerance of DHFR as a function of TYMS background. The heatmap shows the fraction of DHFR mutations with growth rates of 0.9 or better as TYMS *k_cat_* and K_m_ are discretely varied. Both axes are natural log spaced, TYMS *k_cat_* was sampled at 50 points between ln(-2) and ln(2), while TYMS K_m_ was sampled at 50 points between ln(-1) and ln(3). The values for TYMS R166Q, Q33S and WT are marked with red, cyan and black circles respectively. **g)** A zoomed-in version of (**f**), focusing on the mutational tolerance of DHFR for TYMS backgrounds similar in velocity to WT and Q33S TYMS.

Having established model performance on this subset of 114 biochemically characterized DHFR and TYMS sequences, we next examined consistency of the model with all growth rate measurements (the total model fit). However the effect of most mutations on catalysis is unknown. Thus, for each DHFR point mutant we used Monte Carlo sampling to identify a space of *k_cat_* and K_m_ values consistent with the three growth rate measurements (in the three TYMS backgrounds). While three growth rate measurements were insufficient to uniquely constrain both *k_cat_* and K_m_ (the solution space is degenerate), this process did permit estimation of log_10_ catalytic power (*k_cat_*/K_m_) for all 2,696 characterized point mutants. For the subset of biochemically characterized DHFR mutants we observed reasonable agreement between the *in vitro* measurements and those inferred from our experimental growth rate data (R^2^ = 0.74, **Fig. 5c**). Once catalytic parameters were estimated across all point mutants, we put them back into the model to assess the correspondence between the predicted (modeled) growth rates, predicted epistasis, and our experimental observations, yielding a global picture of model fit quality. Overall, we observed that the model well-described the data with two exceptions. First, there was a small proportion of DHFR mutations that were predicted to be rescued by TYMS R166Q but in actuality were not (70 total, 2% of all DHFR mutations, the horizontal stripe of red dots in **Fig. 5d**). It is possible that these mutations caused a growth rate defect through DHFR mis-folding and aggregation, a factor not captured by our model. Second, there was a proportion of DHFR mutations predicted to have negative epistasis to TYMS R166Q but observed to exhibit mild positive epistasis (**Fig. 5e**). These differences may be related to the fact that DHFR abundance is modeled with a single parameter across all mutants. These discrepancies could be addressed in future work by including additional high-throughput assay data on stability or abundance to tease apart mutational effects on catalysis from stability^35^. Nevertheless, the data indicated that our model can globally describe growth rate phenotypes given variation in enzyme velocity.

The resulting model and inferred catalytic parameters now permit estimation of DHFR single mutant fitness in any TYMS background. We computed the fraction of DHFR point mutants that are neutral (growth rate above 0.9) as a function of variation in TYMS *k_cat_* and K_m_. These calculations highlighted that selection on DHFR activity is strongly shaped by TYMS background, with low-activity TYMS variants increasing the mutational tolerance of DHFR (**Fig. 5f,g**). This suggests that TYMS inhibition or loss of function could promote the evolvability of DHFR both in the clinic and laboratory settings.

### Epistasis between DHFR and TYMS is organized into structurally localized groups

Next, we examined the structural pattern of DHFR positions with epistasis to TYMS Q33S and TYMS R166Q. Given that mutations tend to have similar epistatic effects at a particular DHFR position in our data set (**Fig. S4**), we used k-means clustering to sort positions into four categories according to their pattern of epistasis: negative, insignificant, positive, and strong positive (**Fig. 6a, Table S6**). The strong positive category solely contained DHFR mutations in the TYMS R166Q background, while the negative epistasis category was predominantly occupied by DHFR mutations in the TYMS Q33S background. Mapping these positions to the DHFR structure showed that epistasis is organized into spatially distinct regions of the tertiary structure (**Fig. 6b,c**). Mutations with negative epistasis to Q33S tended to be proximal to the DHFR active site, particularly the folate binding pocket. The negative epistasis cluster included several key positions near or in the Met-20 loop, which is known to undergo conformational fluctuations associated with catalysis(residues A9, V13, E17 and M20) ^28,31^. It also encompassed positions I5, L24, L28, and F31 which surround the folate substrate. Several of these positions have known roles in catalysis; mutations at position 31 promoted product release (while slowing hydride transfer), and dynamics of the M20 loop (which includes V13,E17) are associated with substrate binding and product release^32,36^. Additionally, specific mutations at positions 5, 20, and 28 result in trimethoprim resistance by altering trimethoprim affinity^36^. These structural and biochemical observations are consistent with the finding that mutations with negative epistasis tended to yield moderate to severe growth rate defects. In contrast, positions with positive epistasis to Q33S often had very little (or sometimes a beneficial) effect on growth rate, and were distributed around the DHFR surface (**Fig. 4c**, **Fig. 6b**). In the context of TYMS R166Q only one position — C85 — was included in the negative epistasis cluster (**Fig. 6c**). A large fraction of DHFR positions (53%, 84 total) displayed positive epistasis to TYMS R166Q; these positions were distributed throughout the DHFR structure. The positions in the strong positive epistasis cluster included mutations with some of the most severe effects on growth rate in the WT TYMS context. A number of these positions were previously established as important to DHFR catalysis, including residues F31, L54, G121, D122, and S148^28^. Mutations at these sites can be detrimental to *k_cat_*, K_m_, or both.

**Figure 6.**
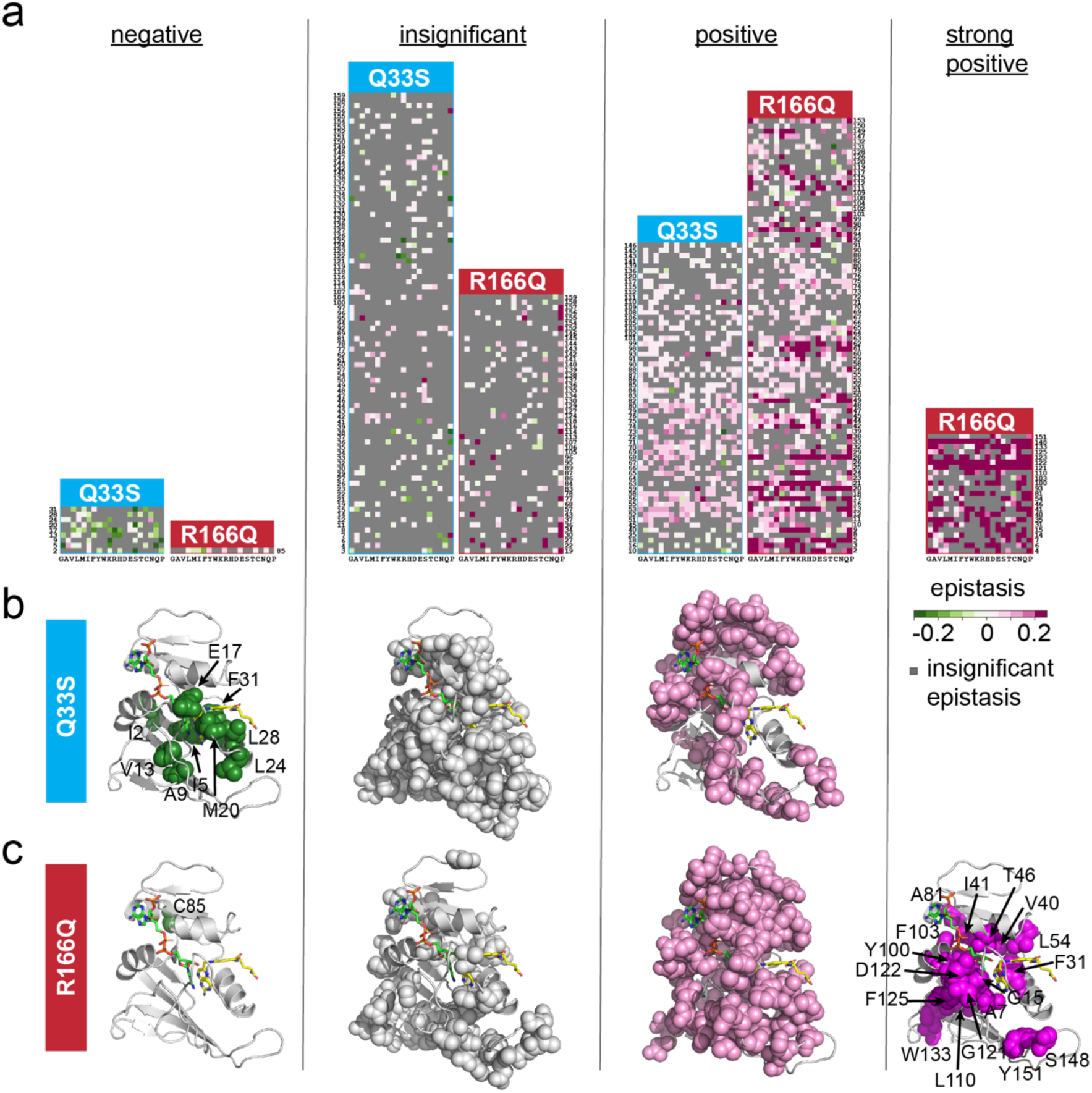
DHFR positions clustered by epistatic mutational effect. **a)** Clusters of DHFR positions organized by predominant epistasis type. In each heat map DHFR positions are ordered along the vertical axis; amino acid residues are organized by physiochemical similarity along the horizontal axis. As in earlier plots, green indicates negative epistasis, and pink indicates positive epistasis. Grey pixels mark mutations with statistically insignificant epistasis. **b)** Structural location of epistatic clusters for DHFR to TYMS Q33S. The DHFR backbone is in grey cartoon (PDBID: 1RX2^31^). Folate, the DHFR substrate is indicated with yellow sticks. The NADP+ cofactor is in green sticks. **c)** Structural location of epistatic clusters for DHFR to TYMS R166Q. Color coding is identical to panel (**b**).

### Epistasis and the structural encoding of DHFR catalysis

When the epistatic clusters are viewed together on the structure, one sees that they form approximate distance-dependent shells around the active site (**Fig. 7a-d**). Considering the pattern of epistasis to TYMS Q33S, positions with negative epistasis were closest to the active site, surrounded by positions with insignificant epistasis, and finally positions in the positive epistasis cluster form an outer shell (**Fig. 7a,b**). For TYMS R166Q, positions in the strong positive epistasis cluster were closest to the active site, followed by positive epistasis positions, and finally those with insignificant epistasis (**Fig. 7c,d**). For comparison, we also mapped the model-predicted catalytic power averaged across all mutations at a position to the structure (**Fig. 7e**). Together, these structural images paint a picture of the molecular encoding of catalysis and epistasis. Mutations with predicted intermediate-to-large effects on catalysis were nestled near the active site and showed negative epistasis to Q33S and strong positive to positive epistasis to R166Q. Mutations with more mild effects on catalysis showed weaker positive to insignificant epistasis to R166Q and Q33S. Though catalysis and epistasis showed an approximate distance-dependent relationship to the DHFR active site, there a number of key positions distal to the active site that exhibited large growth rate effects, strong positive epistasis to TYMS R166Q, and likely act allosterically to tune catalytic activity (e.g. L110, G121, D122, W133, S148, and Y151). The positions with the largest estimated effects on catalysis were highly evolutionarily conserved (P < 10^-10^ by Fisher’s exact test, **Table S7**, **Fig. 7f**), indicating that our model and experimental data are capturing features relevant to the fitness of DHFR. Together, these data show that TYMS metabolic context strongly modulates the constraints on DHFR activity and catalysis.

**Figure 7.**
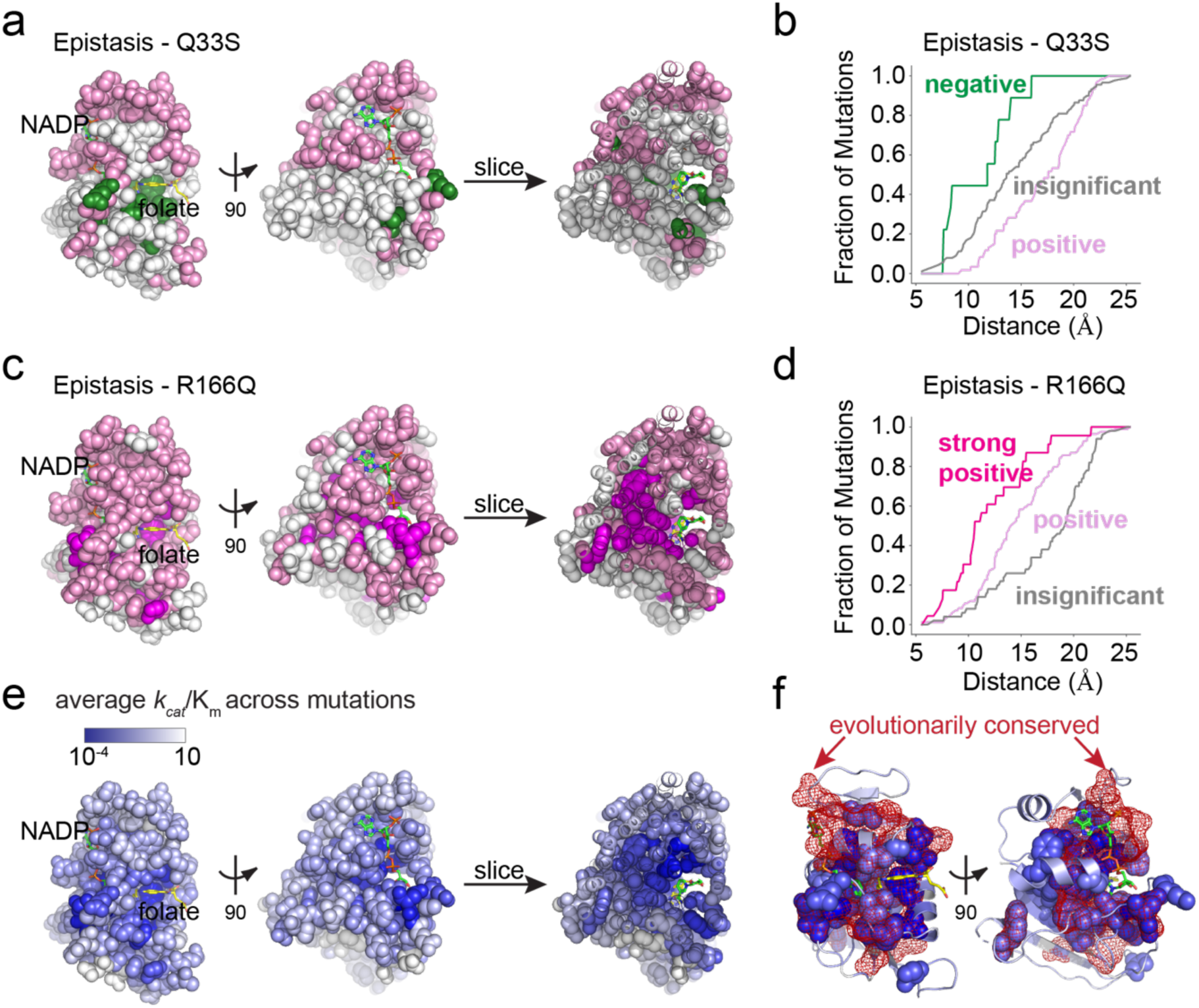
The structural organization of epistasis in DHFR. **a)** Epistasis of individual DHFR positions to TYMS Q33S. The DHFR structure is shown in space filling spheres (PDBID: 1RX2), with the NADP co-factor in green sticks, and folate in yellow sticks. A slice through the structure shows the interior arrangement of epistasis. Positions in the negative epistasis cluster are colored green, positions in the positive epistasis cluster are colored pink. Grey spheres indicate positions in the insignificant epistasis cluster. **b)** Cumulative distribution of positions in each epistatic cluster by distance to the DHFR active site for the TYMS Q33S background. In this case, active site was defined as the C6 atom of the folate substrate. Color coding follows from (**a**) **c)** Epistasis of individual DHFR positions to TYMS R166Q. The DHFR structure is shown in space filling spheres (PDBID: 1RX2), with the NADP co-factor in green sticks, and folate in yellow sticks. A slice through the structure shows the interior arrangement of epistasis. Positions with strong positive epistasis are colored magenta, positions in the positive epistasis cluster are colored pink. Grey spheres indicate positions in the insignificant epistasis cluster. **d)** Cumulative distribution of positions in each epistatic cluster by distance to the DHFR active site for the TYMS R166Q background. In this case, active site was defined as the C6 atom of the folate substrate. Color coding follows from (**c**) **e)** The position-averaged log_10_ catalytic power across measured mutations. All residues are indicated in space filling and color coded by the average mutational effect. Blue indicates positions where mutations have a deleterious effect on catalytic power (on average), while white indicates mutations that have little to no effect on catalytic power. Again, the NADP co-factor is shown in green sticks, and folate in yellow sticks. A slice through the structure shows the interior distribution of mutational effects on catalysis. **f)** The structural overlap between positions associated to catalysis and evolutionary conservation. The DHFR backbone is shown in grey cartoon, the NADP co-factor in green sticks, and folate in yellow sticks. Positions where mutations have (on average) a deleterious effect on catalysis at least half a standard deviation below the mean are shown in blue space filling (color coding identical to **c**). Evolutionarily conserved positions (as computed by the Kullback-Leibler relative entropy in a large alignment of DHFR sequences) are outlined in red mesh.

## DISCUSSION

It is well-appreciated that physical protein interactions place constraints on the individual interacting monomers. Protein interfaces are organized to bind with appropriate affinity and avoid non-specific interactions^37,38^. The individual components of physical complexes tend to be expressed in similar ratios to avoid dosage related toxicity and aggregation^39,40^. However the extent to which interactions mediated by biochemistry (rather than binding) constrain the function and sequence of individual monomers has remained less clear. We have explicitly revealed these interactions at single-residue resolution for one model system and coupled them with a mathematical model to quantify the intracellular constraints on DHFR and TYMS relative catalytic activities.

Our mutagenesis data and modeling show that TYMS activity strongly modifies the constraints on DHFR catalytic parameters; shaping both the range and relative importance of *k_cat_* and K_m_ in modulating growth. This biochemical interaction results in an approximately shell-like pattern of mutational sensitivity to TYMS background (epistasis) in the DHFR tertiary structure. Extreme loss-of-TYMS function buffered variation in some of the most conserved DHFR active site positions, while moderate loss of function buffered variation at more peripheral surface exposed sites. Given these data, we expect that inhibition or loss-of-function in TYMS will promote the evolvability of DHFR in nutrient rich environments, a finding with consequences for both laboratory and clinical evolution. For example, inhibiting TYMS activity in the clinic may promote the evolution of drug resistance in DHFR, while activating TYMS may restrict evolutionary accessible paths. In the laboratory, strains with reduced TYMS activity could provide a less stringent context for testing designed sequences or evolving new DHFR function.

The existence of an enzyme velocity to growth-rate mapping — by definition — allows us to relate variation in DHFR and TYMS catalytic parameters to growth rate. It also allows one (in principle) to do the inverse: infer *in vitro* catalytic parameters from growth rate measurements. The intuition follows from classic steady-state Michaelis Menten experiments: to quantify steady state kinetics *in vitro* one measures enzyme initial velocity as a function of substrate concentration. In our sequencing-based experiments, variation in TYMS background effectively titrates intracellular concentrations of DHF (substrate) while growth rate provides an estimate of velocity. Though our current dataset of three TYMS backgrounds is insufficient to uniquely constrain precise fits for *k_cat_* and K_m_, we anticipate that the addition of a few additional TYMS backgrounds and/or the use of more sophisticated fitting approaches will permit more accurate biochemical parameter inference. Indeed, recent work on the small peptide binding proteins (the PDZ and SH3 domains) has shown how measuring the growth rate effect of mutations in different genetic backgrounds and assay conditions can well-constrain biophysical parameters for binding affinity and protein stability^35,41^. One might follow a conceptually similar strategy to learn quantitative biochemical parameters from high throughput growth rate data. New microfluidics-based approaches for high-throughput biochemistry could play a key role in refining and testing such methodology^42^.

Together our findings shape how we think about designing enzymes and metabolic systems. Typical strategies for designing enzymes do not explicitly consider cellular context^43^. As a result, a significant fraction of designs could fail simply because they are not properly “matched” in terms of velocity to the surrounding pathway. The limited ability of homologs to complement growth in another species has been observed for a number of enzymes^44–48^, including DHFR^49,50^. Thus, even a well-designed catalytically active synthetic enzyme could fail to rescue growth if placed in the wrong cellular context. Just as computational protein design considers entire physical complexes to create binding interactions with altered affinity and specificity, one might consider the joint design of biochemically-interacting enzymes to alter metabolic efficiency and growth. Further study of enzyme rates and abundance across species, as well as characterizations of enzyme velocity to growth rate mappings, will help shape our understanding of the system level constraints placed on metabolic enzymes.

## MATERIALS AND METHODS

### EXPERIMENTAL MODEL AND SUBJECT DETAILS

#### Escherichia coli expression and selection strains

ER2566 Δ*folA* Δ*thyA E. coli* were used for all growth rate measurements; this strain was a kind gift from Dr. Steven Benkovic and is the same used in Reynolds et al., 2011 and Thompson et al., 2020^51,52^. XL1-Blue *E. coli* (genotype: *recA1 endA1 gyrA96 thi-1 hsdR17 supE44 relA1 lac* [F’ *proAB lacI*^q^*ZΔM15* Tn*10*(Tet^r^)]) from Agilent Technologies were used for cloning, mutagenesis, and plasmid propagation. BL21(DE3) *E. coli* (genotype*: fhuA2 [lon] ompT gal (λ DE3) [dcm] ΔhsdS*. *λ DE3 = λ sBamHIo ΔEcoRI-B int::(lacI::PlacUV5::T7 gene1) i21 Δnin5*) from New England Biolabs were used for protein expression.

#### Selection vector for DHFR constructs

DHFR variants were cloned into a modified version of the pACYC-Duet 1 vector (Novagen), which we refer to as pTet-Duet. pTet-Duet is a low-copy number vector containing two multiple cloning sites; the first is under control of the T7 promoter and the second was modified to be regulated by the tetracycline repressor (TetR). DHFR (*folA*) is cloned into the first MCS; TYMS (*thyA*) is cloned into the second MCS. During selections we do not induce expression of either gene but instead rely on leaky expression in ER2566 Δ*folA* Δ*thyA E. coli*. The vector map for these constructs can be found on Addgene: 81596.

#### Expression vector for DHFR constructs

*E.coli folA* (the gene encoding DHFR) was cloned into the pHis8-3 expression vector using restriction sites NcoI and XhoI. DHFR was tagged in-frame with an N-terminal 8X-Histidine tag separated from the *folA* reading frame by a thrombin cleavage site. Individual point mutant clones were constructed using the Quikchange II site-directed mutagenesis kit (Agilent).

#### Expression vector for TYMS constructs

The *thyA* gene (encoding TYMS) was amplified by PCR from *E. coli* MG1655 and cloned into the vector pET24A using XbaI/Xho restriction sites. The point mutants of TYMS (Q33S, R127A, and Q33S) were made using the Agilent QuikChange II site-directed mutagenesis kit.

## METHOD DETAILS

### Plate-reader Based Growth Rate Assays

DHFR and TYMS point mutant combinations in the selection vector were transformed into ER2566 Δ*folA* Δ*thyA* chemically competent cells by heat shock. The cells were recovered for 60 minutes at 37°C with shaking at 220 rpm, spread on agar plates (Luria Broth (LB) containing 30 µg/ml chloramphenicol and 50 µg/ml thymidine), and grown at 37°C overnight. The next day, liquid overnight cultures were inoculated from a streak over multiple colonies and grown overnight at 37°C in LB supplemented with 30 µg/ml chloramphenicol and 50 µg/ml thymidine. These overnight cultures were pelleted and washed with M9 minimal media, then resuspended in pre-warmed M9 media supplemented with 0.4% glucose, 0.2% amicase, 2 mM MgSO_4_, 0.1µM CaCl_2_, 30 µg/ml chloramphenicol (henceforth referred to as M9 selection media). Next, OD_600_ for all resuspended cultures was measured in a Perkin Elmer Victor X3 plate reader. Cultures were then diluted to OD_600_=0.1 in prewarmed M9 selection media with 50 µg/ml thymidine and incubated for 4 hours at 30°C, shaking at 220 rpm. After this period of adaptation and regrowth, cultures were back-diluted to OD_600_ = 0.1 in 1 ml prewarmed M9 selection media with 50 µg/ml thymidine. These cells were inoculated into 96-well culture plate at OD_600_ = 0.005 (10 µl cells into 200 µl total well volume) containing prewarmed M9 selection media with 50 µg/ml thymidine; plates were sealed with EasySeal permeable covers (Sigma Aldrich). All growth rate measurements were made in triplicate. Plates were shaken for 10 seconds before reading, and Readings of OD_600_ were taken every 6 minutes over 24 hours using a BioTek Synergy Neo2 plate reader in a 30°C climate-controlled room.

### DHFR Saturation Mutagenesis Library Construction

The DHFR saturation mutagenesis library was constructed as four sub-libraries in the pTet-Duet selection vector (see above for selection vector details). Each sub-library combines mutations within 40 contiguous amino acid positions to ensure that the mutated region can be completely covered with short read sequencing (a 300 cycle v2 Illumina sequencing kit). The regions spanned by each sub-library were as follows: amino acid positions 1-40 (sub-library 1, SL1), 41-80 (sub-library 2, SL2), 81-120 (sub-library 3, SL3), and 121-159 (sub-library 4, SL4). ‘Round the Horn’ or inverse PCR (iPCR) with mutagenic NNS primers (N = A/T/G/C, S = G/C) was used to introduce all 20 amino acid substitutions at a single amino acid position as described in Thompson et al^52^. Library completeness was verified by deep sequencing. In our initial validation sequencing run we found that mutations at positions W22 and L104 were systematically under-represented; iPCR was repeated for these positions and they were supplemented into their respective assembled sub-libraries.

After sub-library assembly, restriction digest and ligation were used to subclone each sublibrary into pTet-Duet plasmids containing the three different TYMS backgrounds (WT, R166Q, or Q33S). The entire DHFR coding region containing restriction sites (NotI and EcoNI) was amplified by PCR. PCR reaction was size-verified with agarose gel electrophoresis with an expected band size of 627 bp. The library PCR products and target plasmids were double digested with NotI and EcoNI for 3 hours at 37°C. To prevent re-circularization, the digested plasmid was treated with Antarctic phosphatase for 1 hour at 37°C. The DHFR insert and treated plasmid were ligated with T4 DNA ligase overnight at 16°C. The concentrated ligation product was then transformed into *E. coli* XL1-blue by electroporation, and recovered in SOB for 1 hour at 37°C. 20 µL of the recovery culture was serially diluted and plated on LB-agar with 50 µg/mL thymidine and 30 µg/mL chloramphenicol, to permit quantification of transformants and estimate library coverage following ligation into the alternate TYMS backgrounds. The minimum library coverage was 1000 colony forming units (CFU) per mutation. The remaining recovery culture was grown in a flask containing 12 ml LB with 30 µg/mL chloramphenicol and 50 µg/mL thymidine at 37°C, with 220 rpm shaking overnight. 10 ml of the overnight culture was miniprepped with the Gene-Jet Mini-prep kit (Fisher Scientific, K0503) to obtain the plasmid library.

### Growth Rate Measurements in the Turbidostat for all DHFR mutant libraries

All sublibraries were inoculated, grown, and sampled in triplicate. Each plasmid sub-library was transformed into the *E. coli* double knockout strain ER2566 Δ*folA* Δ*thyA* by electroporation and recovered in SOB for one hour at 37°C. At this transformation step we again estimated library coverage to ensure the complete library was transformed into our selection strain. To estimate library coverage, 20 µL of the recovery culture was serial diluted with SOB and plated on LB agar plates containing 30 µg/mL chloramphenicol and 50 µg/mL of thymidine. The remainder of the recovery culture was inoculated into M9 selection media supplemented with 50 µg/mL thymidine and grown overnight at 37°C. All selection experiments in this work had an estimated library coverage of 1000 CFU/mutant or greater. The overnight liquid culture was washed and back-diluted to OD_600_=0.1 in M9 selection media supplemented with 50 µg/mL thymidine, and incubated for four hours at 30°C to allow adaptation to selection temperature and to return the culture to log-phase growth. Following adaptation, selection was initiated by back-diluting these cultures to an OD_600_ of 0.1 into 17 mL of pre-warmed M9 selection media supplemented with 50 µg/mL thymidine in continuous culture vials. These vials were then incubated in a turbidostat with a target OD_600_ of 0.15 at a temperature of 30°C. The turbidostat maintained a set optical density by adding 2.8 mL dilutions of M9 selection media supplemented with 50 µg/mL thymidine in response OD detection, and was built according to the design of Toprak et al^53^. Culture samples (1 mL each) were taken at the beginning of selection (t = 0 hours) and at 4, 8, 12, 20, and 24 hours into selection. Immediately after each time point, these 1 mL samples were pelleted by centrifugation, supernatant removed, and stored at −20°C.

### Next Generation Sequencing Amplicon Sample Preparation

Each turbidostat selection sample (representing a particular timepoint for a sub-library and replicate) was prepared for sequencing as a PCR amplicon using Illumina TruSeq-HT i5 and i7 indexing barcodes. To generate these amplicons, each cell pellet from the growth rate assay was thawed and lysed by resuspending the cells with 100 µL dH_2_O and incubation at 95°C for 5 minutes. Lysates were then clarified by centrifugation at maximum speed for 10 minutes in a room temperature bench top microcentrifuge. Supernatants containing plasmids were isolated from the pellet. 1 µL of each supernatant was used as the template for an initial round of PCR with Q5-Hot Start Polymerase (NEB) that amplified the DHFR coding region of the sublibrary (10 PCR cycles total, standard Q5 reaction conditions). From this first PCR reaction, 1 µL was used in a second round of PCR (22 cycles of denaturation/anneal/elongation) with primers that added Illumina sequencing adaptors. Together, these two rounds of PCR yielded a final product of size: 315 bp (SL1), 308 bp (SL2), 298 bp (SL3), 304 bp (SL4). The amplicons were size verified using 1% agarose gel electrophoresis. In the case where a sample did not produce an amplicon, the first round PCR was repeated with 2 µL of the supernatant rather than 1 µL, with the remaining preparation identical. All amplicons were individually quantified using with Quant-iT™ PicoGreen™ dsDNA Assay Kit (ThermoFischer Scientific) and mixed in equimolar ratio, with a final target amount greater than or equal to 2000 ng. Errors in pipetting volume were minimized by ensuring that more than 2 µL was taken from each amplicon. This mixture was gel-purified and then cleaned and concentrated using the Zymo Research DNA Clean & Concentrator-5 kit. Purity was assessed by A260/A80 and A260/A230 nm absorbance ratios. The sample library DNA concentration was measured using a Qubit dsDNA HS Assay in a Qubit 3 Fluorometer (Invitrogen by Thermo Fischer Scientific). The sample library was diluted to 30 nM in a volume of 50 µL of TE buffer (1 mM Tris-HCl (pH 8.5), 10 mM EDTA (pH 4)). This mixed and quantified library was sequenced on an Illumina HiSeq (150 cycle x 2 paired-end) by GeneWiz. The NGS sequencing run resulted in 252 GB of data, consisting of 337,353,664 reads. The raw data have been deposited with the NCBI sequencing read archive under project identifier PRJNA791680.

### DHFR Expression and Purification

DHFR mutant variants were expressed in BL21(DE3) *E. coli* grown at 30°C in 50 ml Terrific Broth (TB) with 35 µg/ml Kanamycin (Kan) for selection. Expression was induced at an OD_600_ = 0.6-0.8 with 250 uM IPTG, and cells were grown at 18°C for 16-18 hours. Cultures were pelleted by centrifugation for 10 minutes at 5000 x g, 4°C and supernatant removed; cell pellets were stored at −80°C. Thawed cell pellets were lysed by sonication in 10ml lysis buffer (50 mM Tris, 500 mM NaCl, 10 mM imidazole, pH 8.0 buffer containing 0.1 mM PMSF, 0.001 mg/ml pepstatin, 0.01 mg/ml leupeptin, 20 µg/ml DNAseI and 5 µg/mL lysozyme). The resulting lysate was clarified by centrifugation and incubated with 0.1ml Ni-NTA agarose (Qiagen) slurry (0.05 ml column volume) equilibrated in Nickel Binding Buffer (NiBB, 50 mM Tris pH 8.0, 500 mM NaCl, 10 mM imidazole) for 15 minutes on a tube rocker at 4°C. The slurry was then transferred to a disposable polypropylene column (BioRad). After washing with 10 column volumes (CV) of NiBB, DHFR was eluted with 0.5 mL 50mM Tris pH 8.0, 500 mM NaCl, 400 mM imidazole. The eluted protein was concentrated and buffer-exchanged to 50 mM Tris, pH 8.0 in a 10 kDa Amicon centrifugal concentrator (Millipore) and centrifuged 15 min at 21,000 x g, 4°C to pellet any precipitates. Following buffer exchange, the protein was purified by anion exchange chromatography (using a BioRad HiTrapQ HP column on a BioRad NGC Quest FPLC). A linear gradient was run from 0-1 M NaCl in 50 mM Tris pH 8.0 over 30 ml (30 column volumes, CV) while collecting 0.5 ml fractions. Fractions containing DHFR were combined, concentrated, flash-frozen in liquid nitrogen, and stored at −80°C.

### TYMS Expression and Purification

Individual TYMS mutants were expressed in BL21(DE3) *E. coli* grown at 37°C in 50 ml Terrific Broth (TB) with 35 µg/ml Kanamycin (Kan) for selection. Expression was induced with 1mM IPTG when the cells reached an OD_600_ = 0.6-0.8, and the cells were then grown at 18°C for 16-18 hours before harvesting pellets for storage at −80°C. TYMS was purified from the frozen pellets following a protocol adapted from Changchien et al ^54^. Cell pellets were thawed and resuspended in TYMS lysis buffer (20 mM Tris, 10 mM MgCl2, 0.1% deoxycholic acid, pH 7.5 with 5 mM DTT, 0.2 mg/ml lysozyme, 5 µg/ml DNAse I) and incubated at room temperature while rocking for 15 minutes. The resulting supernatant was clarified by centrifugation. Next, streptomycin sulfate was added to a final concentration of 0.75% to separate nucleic acids. The cells were incubated rocking at 4°C for 10 minutes and the supernatant was retained following centrifugation for 10 minutes at >10,000 x g. Ammonium sulfate was then added at 50% saturation (0.3 g/ml), mixed for 10 minutes at 4°C, then centrifuged as above, retaining supernatant. Additional ammonium sulfate was then added to the supernatant at 80% saturation (an additional 0.2g/ml), mixed for 10 minutes at 4°C, and centrifuged as above, retaining the pellet. The pellet was dissolved in 25mM potassium phosphate pH 6.5 and dialyzed overnight at 4°C against 1L 25 mM potassium phosphate pH 6.5. Following dialysis the protein was purified by anion exchange (HiTrap Q HP column, Cytiva) with a 25 CV linear gradient from 0M NaCl to 1M NaCl in 25mM potassium phosphate pH 6.5. FPLC fractions containing TYMS were combined and concentrated using a 10 kDa Amicon concentrator (Millipore) and stored at 4°C for up to a week.

### DHFR Steady-state Michaelis Menten Kinetics

DHFR *k_cat_* and K_m_ were determined under Michaelis-Menten conditions with saturating concentrations of NADPH as in prior work^55,56^. Briefly, DHFR protein concentration was determined by measuring A_280_ (extinction coefficient = 33500 M^-1^cm^-1^). DHF (Sigma Aldrich) was prepared in MTEN buffer (50 mM MES, 25 mM Tris base, 25 mM Ethanolamine, 100 mM NaCl, pH 7.0) containing 5 mM DTT (Sigma Aldrich). 100 nM DHFR protein and 100 µM NADPH (Sigma Aldrich) were combined in MTEN buffer with 5 mM DTT and pre-incubated for 1 hour at 25°C prior to measurement. To initiate the reaction, the protein-NADPH solution was mixed with DHF in a quartz cuvette (sampling DHF over a range of concentrations, tuned to the Km of the mutant). The initial velocity of DHFR was measured spectrophotometrically by monitoring the consumption of NADPH and DHF (decrease in absorbance at 340 nm, 1¢c_340_=13.2 mM^-1^ cm^-1^). All measurements were made in triplicate; analysis was performed using the Michaelis-Menten nonlinear regression function of Graph Pad Prism.

#### Preparation of TYMS substrate for assaying enzyme activity and steady state kinetics

(6R)-methylenetetrahydrofolic acid (MTHF, CH_2_H_4_fol) was purchased from Merck & Cie (Switzerland) and dissolved to 100 mM in nitrogen-sparged citrate-ascorbate buffer (10 mM ascorbic acid, 8.5 mM sodium citrate, pH 8.0). 30 µL aliquots were made in light-safe microcentrifuge tubes, flash-frozen in liquid nitrogen, and stored at −80C. Before use, the stock was thawed and diluted to 10 mM in TYMS kinetics reaction buffer (100 mM Tris base, 5 mM Formaldehyde, 1 mM EDTA, pH 7.5) and quantified in an enzymatic assay: 50 µM MTHF, 200 µM dUMP and 1µM TYMS protein were combined and A_340_ measured until steady-state reached. Actual concentration was then calculated from the difference in A_340_ before and after the reaction using Beer’s Law (MTHF extinction coefficient: 6.4 mM^-1^cm^-1^).

### TYMS Steady-state Michaelis Menten Kinetics

TYMS *k_cat_* and K_m_ were determined for both dUMP and MTHF under Michaelis-Menten conditions by varying one substrate and holding the other saturating as in prior work^57,58^. Briefly, TYMS protein concentration was determined by measuring A_280_ (extinction coefficient = 53400 M^-1^cm^-1^). TYMS protein was prepared in TYMS assay buffer (100 mM Tris base, 5 mM Formaldehyde, 1 mM EDTA, pH 7.5) containing 50 mM DTT (Sigma Aldrich). 50 nM TYMS protein and either 100 µM dUMP (Sigma Aldrich) or 150 µM MTHF (Merck & Cie) were combined with varying concentrations of the other substrate to initiate the reaction. The production of DHF was monitored spectrophotometrically (increase in absorbance at 340nm, 1¢E_340_=6.4 mM^-1^cm^-1^) for 2 minutes per reaction. All measurements were made in triplicate; analysis was performed using the Michaelis-Menten nonlinear regression function of Graph Pad Prism.

## QUANTIFICATION AND STATISTICAL ANALYSIS

### Enzyme Velocity to Growth Rate Model Construction and Parameterization

For the purposes of modeling, we approximated DHFR and TYMS as a two-enzyme cycle in which DHFR produces THF and consumes DHF, and TYMS produces DHF and consumes THF. This abstraction ignores the different carbon-carrying THF species, instead collapsing them into a single “reduced folate” pool. This simplification allows us to construct an analytically solvable model for steady state THF concentration that we can then relate to growth (Equation 3).

First, we fit the free parameters in the Goldbeter-Koshland equation ([DHFR], [TYMS_WT_], [TYMS_R166Q_], fol_tot_) using a set of ten metabolomics measurements for the relative abundance of 3-glutamate form of formyl THF as obtained in prior work^1^. These measurements were made for DHFR mutations G121V, F31Y/L54I, M42F/G121V, F31Y/G121V and the WT in the background of WT TYMS and TYMS R166Q. So why this particular folate species? We noticed that the relative abundance of many of the reduced THF species in our data set was correlated, and chose formyl THF to model because the experimental data were less variable and showed a strong, monotonic relationship with cell growth. Then, we fit the free parameters in equation two (gmax, gmin, K, n) using a set of ten growth rate measurements for the same DHFR/TYMS mutation pairs. This fitting process gave rise to the fits shown in figure 1. When assessing model performance against the larger set of TYMS variants (as in figure 2) we refit all parameters (gmax, gmin, K, n, [DHFR], [TYMS_WT_], [TYMS_R166Q_], [TYMS_Q33S_], [TYMS_R127A_], fol_tot_) to the growth rate data only since we did not have metabolomics data for this larger set. All parameter fits were made in python using the least_squares fitting function of the scipy.optimize module^59^; the complete fitting process is documented in Jupyter notebook 1_KGmodel.ipynb in the associated github repository.

We assessed the model sensitivity to shuffling the data (Figure 2 – supplement 2) by randomly shuffling all catalytic parameters (*k_cat_*, K_m_) 50 times across DHFR and TYMS and computing an R^2^ value. We also assessed model sensitivity to subsampling the data; error bars in Figures 1, 2, and Figure 2 – supplement 2 correspond to SEM across “jackknife” re-samplings of the data wherein one DHFR/TYMS combination was left out for each re-sampling. Finally, to assess the global model fit to the data (as in Figure 6 and Figure 6 – supplement 1) we first fit the 9 model parameters (gmax, gmin, K, n, [DHFR], [TYMS_WT_], [TYMS_R166Q_], [TYMS_Q33S_], fol_tot_) using the growth rate measurements of 16 DHFR mutations for which experimental *k_cat_* and K_m_ were known (48 total observations given the three TYMS backgrounds). Then, fixing these parameters, we fit *k_cat_* and K_m_ values for all 2696 mutations with growth rate measurements in all three TYMS backgrounds to the complete data set of 8,088 sequencing-based growth rate observations. This process is documented in Jupyter notebook 4_ModelAndDMSData.ipynb in the associated github repository.

### Next Generation Sequencing Data Processing and Read Counting

All PCR amplicons (corresponding to individual replicates, timepoints and sublibraries) were sequenced on an Illumina HiSeq using 2 x 150 paired end reads. The resulting fastq files were processed and filtered prior to read counting. Briefly, the forward and reverse reads were merged using USEARCH. Each read was quality score filtered (Q-Score ≥ 20) and identified as a WT or mutant of DHFR using a custom python script. This python script filtered for full length reads and base call quality scores greater than 20 (error rate ≤ 1:100). The reads passing these quality control criteria were compared against the wild-type reference sequence to determine mutation identify. Reads that contained multiple point mutations or mutations outside the sublibrary of interest were removed from analysis. This process resulted in counts for the WT and each mutant at each time point and replicate. These counts were further corrected given the expected error in the data (q-score) and Hamming distance from the WT codon to account for potential hopping of WT reads to mutations; a process that was detailed in McCormick et al ^56^.

### Relative Growth Rate Calculations

We calculated relative growth rates for individual mutations and the WT over time from the sequencing-based counts (*N^mut^_t_*, *N^WT^_t_*). Mutants with fewer than 10 counts were considered absent from the data set and were set to zero to reduce noise. From these thresholded counts, we calculated a log normalized relative frequency of each mutation over time:

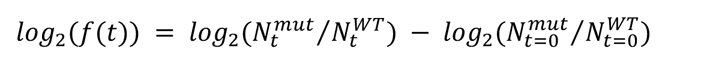

We then calculated relative growth rate (*m*^*TS*^_DHmut_) as the slope of the log relative frequency over time by linear regression. Linear regression was performed using scikit Learn, and individual points were weighted by the number of counts (in order to down weight less-sampled mutants at later time points). Relative growth rate or a mutant was only calculated if the mutant was present over at least the first three time points, otherwise it was classified as a “Null” mutant. Finally, all relative growth rates were normalized such that WT has a relative growth rate of 1. Growth rates were additionally normalized by the bulk culture growth rate (estimated from the turbidostat, in units of generations per hour) to account for small vial-to-vial variations culture doublings across the experiment. The standard error in growth rate was computed across triplicate measurements.

All calculations are shown in Jupyter notebook 2_DMSGrowthRates.ipynb in the associated github repository.

### Epistasis Analysis

Epistasis was calculated according to an additive model:

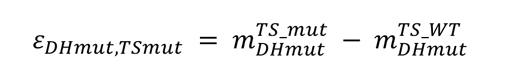

In our experiments TYMS R127A, Q33S and R166Q have no growth rate effect in the WT DHFR context due to thymidine supplementation. Under this formalism, mutations that show improved growth in the mutated TYMS background have positive epistasis, while mutations with reduced growth in the mutated TYMS background have negative epistasis. We assessed the statistical significance of epistasis by unequal variance t-test under the null hypothesis that the mutations have equal mean growth rates in both TYMS backgrounds (across three replicate measurements). These p-values were compared to a multiple-hypothesis testing adjusted p-value determined by Sequential Goodness of Fit (P = 0.035 for TYMS Q33S and P = 0.029 for TYMS R166Q) ^34^. K-means clustering of epistatic positions was performed using a custom script based on that described in Thompson et al ^52^. All epistasis calculations are shown in Jupyter notebook 3_Epistasis.ipynb in the associated github repository.

## ACKNOWLEDGEMENTS

The authors thank Olivier Rivoire for early conversations regarding our data, and Elliott Ross for critical feedback on both the modeling and manuscript. We also thank the Reynolds lab for feedback on experimental design, data analysis, and the manuscript.

## AUTHOR CONTRIBUTIONS

TNN and KAR conceptualized the work and designed experiments. KAR created the mathematical model. TNN collected all deep mutational scanning data. TNN and KAR analyzed the data. CI collected plate-reader based growth rate measurements (used in model development) and performed all Michaelis-Menten enzyme kinetics assays. SMT constructed the deep mutational scanning library and contributed to data interpretation. KAR wrote the original draft. TNN, CI, SMT and KAR revised the paper. KAR provided supervision and funding acquisition.

## DATA AVAILABILITY

The raw sequencing data is available in FASTQ format through the NCBI sequencing read archive, under **BioProject ID PRJNA791680**.

## CODE AVAILABILITY

Code for the enzyme velocity to growth rate model, and analysis of all deep mutational scanning data is available on github: https://github.com/reynoldsk/dhfr-tyms-epistasis

## MATERIALS AND CORRESPONDENCE

The DHFR deep mutational scanning libraries (in all three TYMS backgrounds) have been deposited at addgene under **deposit number 81596**.

Requests for any other materials or data should be made by email to Dr. Kimberly Reynolds, with Christine Ingle in carbon copy (cc).

