## Supplemental Figures and Tables for "The Genetic Landscape of a Metabolic Interaction"

**Contents**:

**I. Supplemental Figures**

**Figure 1**: Growth rates, epistasis, and analysis of model significance for a focused set of mutants.

**Figure 2**: Mutational scanning library completeness.

**Figure 3**: Reproducibility of relative growth rate measurements.

**Figure 4**: Epistasis of DHFR mutations to TYMS background.

**II. Supplemental Tables**

**Table 1:** Steady-state DHFR kinetics parameters.

**Table 2:** Steady-state TYMS kinetics parameters.

**Table 3**: Model parameter fits.

**Table 4:** Relative Growth Rates (and error) for all DHFR mutations in each TYMS background. *(provided as a separate excel file due to size)*

**Table 5**: Epistasis (and p-values) for all DHFR mutations in the TYMS Q33S and R166Q backgrounds. *(provided as a separate excel file due to size)*

**Table 6**: Epistatic residue groups determined by k-means clustering.

**Table 7**: Statistical association between evolutionary conservation and DHFR positions associated to catalysis

**III. Supplemental References**

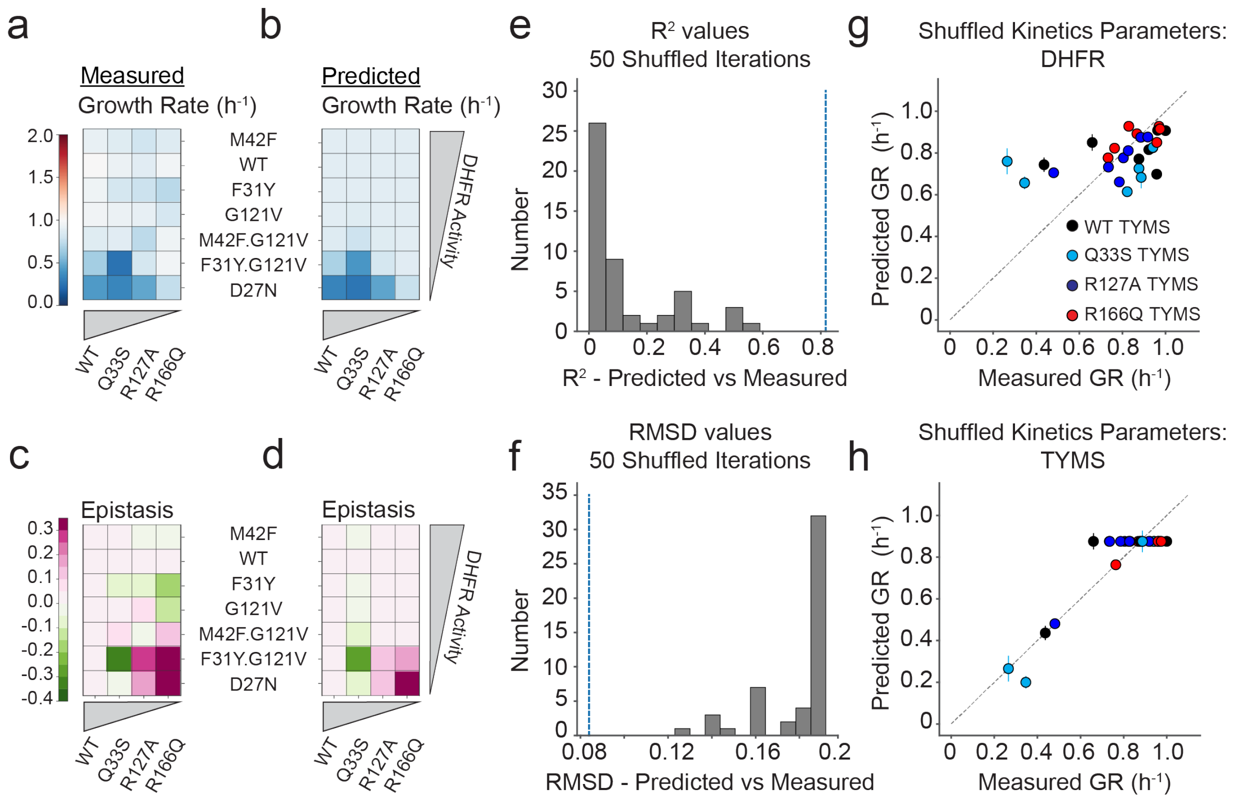


**Supplemental Figure 1**: Growth rates, epistasis, and analysis of model significance for a focused set of mutants.

**a.** Heatmap of experimentally measured growth rates for seven DHFR point mutants in four TYMS backgrounds. White indicates WT-like growth, blue indicates deleterious growth rates.

**b.** Heatmap of computationally predicted growth rates using the best fit model for the data in (**a**).

**c.** Heatmap of experimentally measured epistasis for the same set of DHFR and TYMS mutant combinations. Values were calculated from the data in (**a**). White indicates no epistasis, green indicates negative (amplifying) epistasis, and pink indicates positive (buffering) epistasis.

**d.** Heatmap of computationally predicted epistasis using the growth rates determined in (**b**).

**e.** R-squared values describing the goodness of fit for 50 models trained using randomly shuffled kinetics parameters (shuffling $k_{cat}$ and $K_m$ across all DHFR and TYMS variants). The R-squared value for the true (non-shuffled) data is indicated with a dashed blue line.

**f.** RMSD values describing the deviation between the model and data for 50 models trained using randomly shuffled kinetics parameters (as in **e**). The RMSD value for the true (non-shuffled) data is indicated with a dashed blue line.

**g.** Example model predictions for one particular instance of shuffling DHFR kinetic parameters while retaining the experimentally measured TYMS kinetics parameters. Each point describes a DHFR/TYMS mutant combination, points are color-coded by TYMS background. Error bars in the x axis represent SEM across 3 replicate measurements, error bars on the y axis were obtained by jackknife resampling and refitting the data. The dotted black line marks y=x.

**h.** Example model predictions for one particular instance of shuffling TYMS kinetic parameters while retaining the experimentally measured DHFR kinetics parameters. Error bars in the x axis represent SEM across 3 replicate measurements, error bars on the y axis were obtained by jackknife resampling and refitting the data. The dotted black line marks y=x.

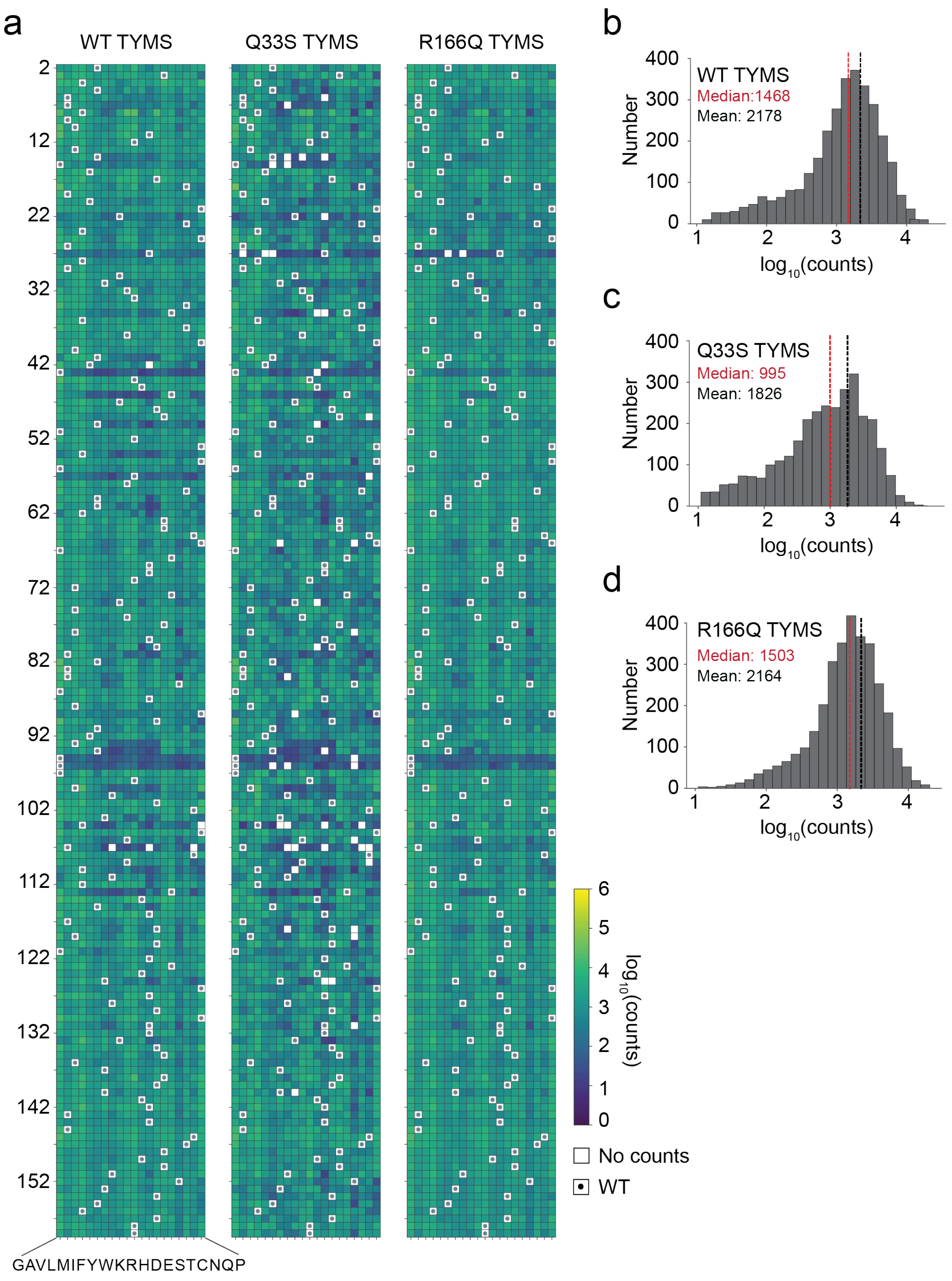


**Supplemental Figure 2:** Mutational scanning library completeness.

a. Heatmaps of $\log_{10}$ sequencing counts (reads) for all single mutations of DHFR in each TYMS background at the start of the experiment (t=0). Representative data are shown for one of three replicate measurements. DHFR positions are indicated along the vertical axis, while amino acids (sorted by physiochemical similarity) are along the horizontal axis. White boxes indicate missing mutations (no counts) and small dots indicate the WT residue identity.

**b.** Distribution of sequencing counts (reads) for all single mutations of DHFR in the WT TYMS background. Median and mean reads are indicated in red and black respectively. Representative data are shown for one of three replicate measurements (same replicate as in A).

**c.** Distribution of sequencing counts (reads) for all single mutations of DHFR in the Q33S TYMS background. Median and mean reads are indicated in red and black respectively. Representative data are shown for one of three replicate measurements (same replicate as in A).

**d.** Distribution of sequencing counts (reads) for all single mutations of DHFR in the R166Q TYMS background. Median and mean reads are indicated in red and black respectively. Representative data are shown for one of three replicate measurements (same replicate as in A).

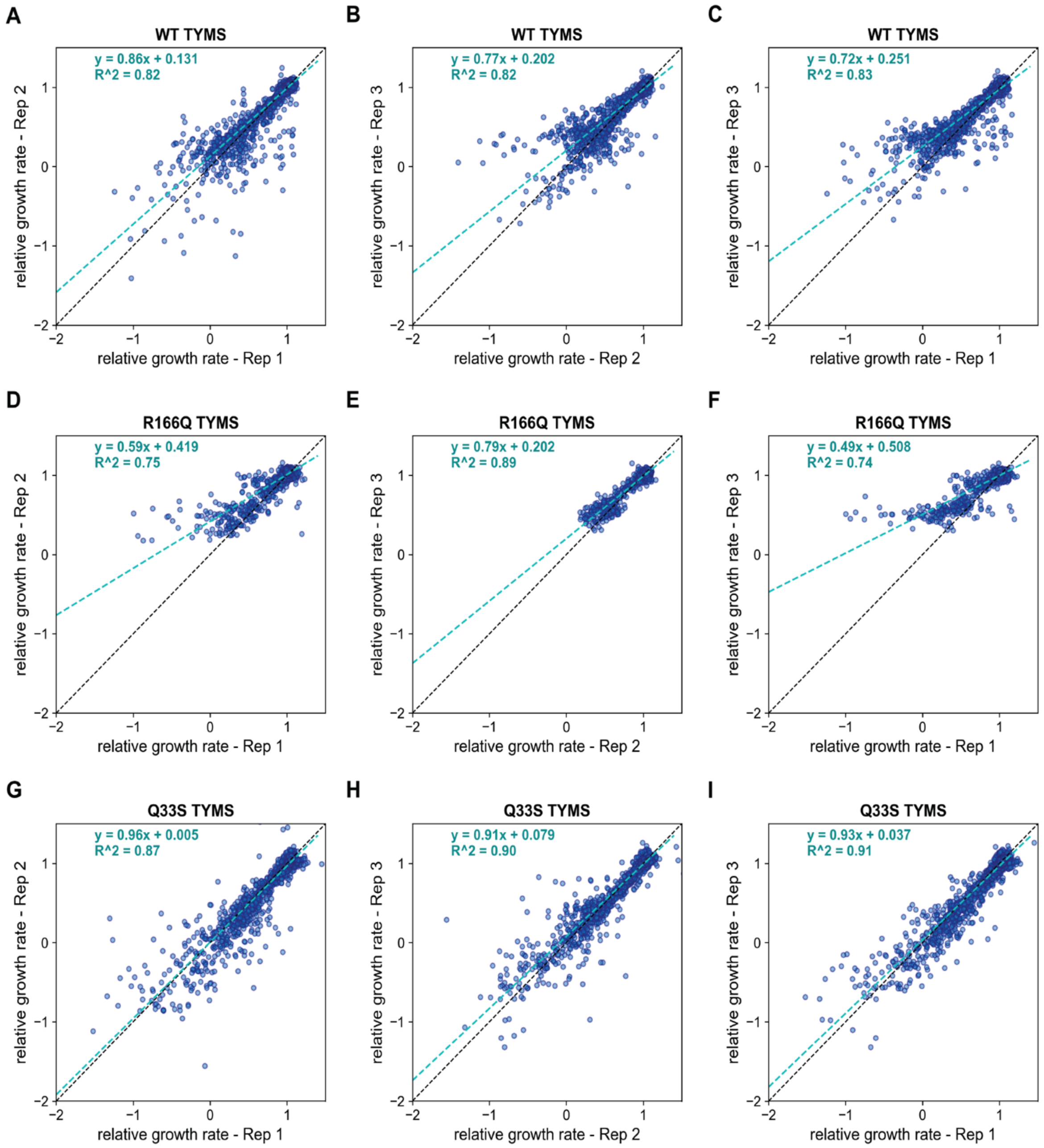


**Supplemental Figure 3:** Reproducibility of relative growth rate measurements.

**a-c)** Correlation in relative growth rate measurements for individual DHFR mutations across three experimental replicates in the WT TYMS background. The cyan dashed line is the line of best fit; black dashed line marks x=y. The data were normalized such that WT growth is one.

**d-f**) Same as A-C, but for the R166Q TYMS background

**g-i**) Same as A-C, but for the Q33S TYMS background

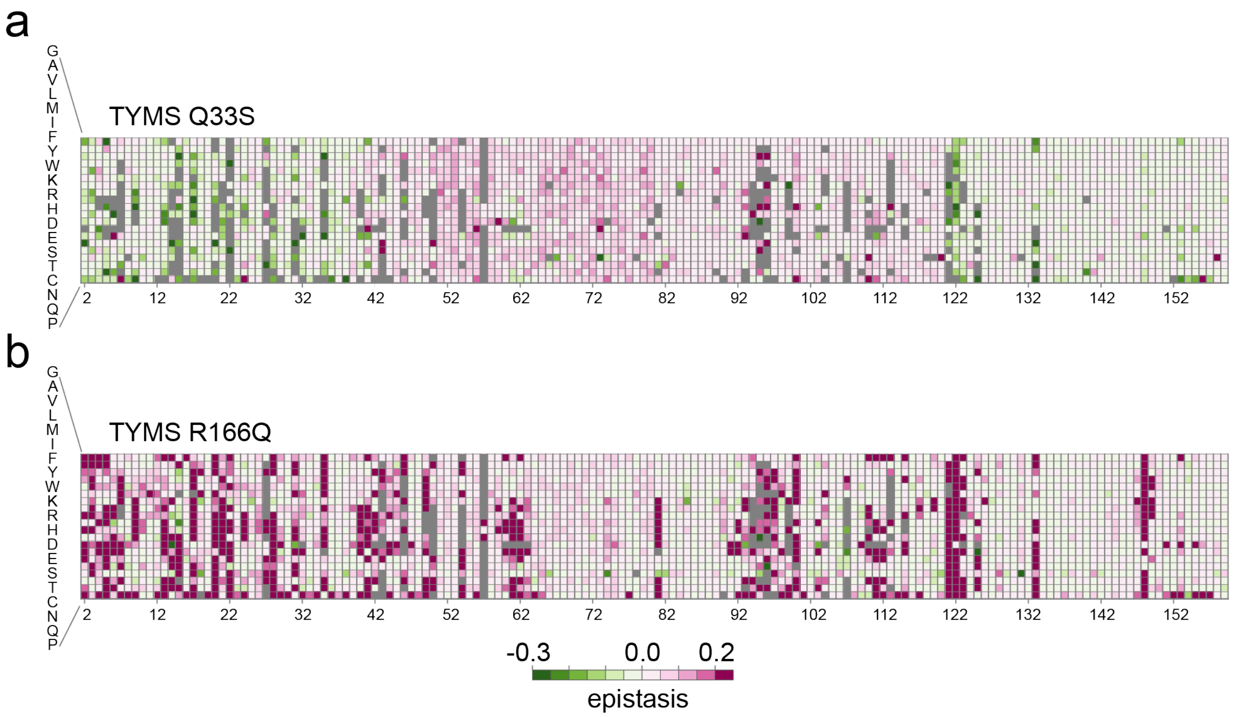


**Supplemental Figure 4**: Epistasis of DHFR mutations to TYMS background

**a.** A heatmap of epistasis for all DHFR positions to TYMS Q33S. Values displayed represent the average across three replicates. Grey pixels indicate mutations that did not have three replicates in both the WT and Q33S TYMS backgrounds. Amino acid mutations are arranged by physiochemical similariy (along the rows), positions are indicated along the columns.

**b.** A heatmap of epistasis for all DHFR positions to TYMS R166Q. Grey pixels indicate mutations that did not have three replicates in both the WT and R166Q TYMS backgrounds. Amino acid mutations are arranged by physiochemical similariy (along the rows), positions are indicated along the columns.

| DHFR mutation | DHFR position | kcat (s-1) | Km (μM) | kcat_std | Km_std | Reference |
|---|---|---|---|---|---|---|
| I5K | 5 | 2.10 | 62.00 | 0.32 | 1.40 | This work |
| I5F | 5 | 3.78 | 7.60 | 0.54 | 1.04 | Tamer et al. MBE 2019 |
| V13H | 13 | 0.70 | 1.90 | 0.05 | 0.10 | This work |
| E17V | 17 | 0.40 | 1.20 | 0.10 | 0.58 | This work |
| M20Q | 20 | 2.85 | 3.00 | 0.68 | 0.57 | This work |
| M20I | 20 | 7.69 | 3.57 | 1.21 | 0.19 | Tamer et al. MBE 2019 |
| P21L | 21 | 3.19 | 4.08 | 0.56 | 0.49 | Tamer et al. MBE 2019 |
| W22H | 22 | 1.89 | 18.00 | 0.06 | 1.20 | Reynolds et al. Cell 2011 |
| A26T | 26 | 3.70 | 7.65 | 0.30 | 1.10 | Tamer et al. MBE 2019 |
| D27N | 27 | 0.05 | 330.00 | 0.01 | 78.90 | Reynolds et al. Cell 2011 |
| D27E | 27 | 14.31 | 56.40 | 1.49 | 5.25 | Tamer et al. MBE 2019 |
| L28F | 28 | 18.50 | 9.90 | 0.62 | 0.80 | Thompson et al. Elife 2020 |
| L28Y | 28 | 19.20 | 21.20 | 0.75 | 1.60 | Thompson et al. Elife 2020 |
| L28R | 28 | 1.13 | 0.95 | 0.05 | 0.09 | Tamer et al. MBE 2019 |
| W30G | 30 | 8.18 | 9.49 | 0.70 | 2.12 | Tamer et al. MBE 2019 |
| W30R | 30 | 8.62 | 4.97 | 1.05 | 0.30 | Tamer et al. MBE 2019 |
| F31V | 31 | 8.65 | 108.00 | 0.29 | 6.80 | Reynolds et al. Cell 2011 |
| F31Y | 31 | 20.61 | 80.00 | 2.12 | 14.00 | Reynolds et al. Cell 2011 |
| F31Y.G121V | 31.121 | 0.13 | 90.60 | 0.01 | 7.40 | Reynolds et al. Cell 2011 |
| F31Y.L54I | 31.54 | 1.94 | 168.30 | 0.16 | 21.40 | Reynolds et al. Cell 2011 |
| V40A | 40 | 24.05 | 7.47 | | | Bershtein et al. PNAS 2012 |
| M42F | 42 | 79.20 | 13.00 | 1.72 | 1.00 | Reynolds et al. Cell 2011 |
| M42F.G121V | 42.121 | 0.40 | 71.80 | 0.04 | 13.20 | Reynolds et al. Cell 2011 |
| L54I | 54 | 7.88 | 35.00 | 0.28 | 3.40 | Reynolds et al. Cell 2011 |
| L54F | 54 | 6.30 | 0.70 | 0.10 | 0.10 | Huang et al. Biochemistry 1994 |
| I61V | 61 | 8.99 | 2.43 | | | Bershtein et al. PNAS 2012 |
| V75H | 75 | 15.50 | 3.39 | | | Bershtein et al. PNAS 2012 |
| V75I | 75 | 18.25 | 3.31 | | | Bershtein et al. PNAS 2012 |
| V88I | 88 | 13.80 | 4.42 | | | Bershtein et al. PNAS 2012 |
| I91V | 91 | 17.87 | 3.48 | | | Bershtein et al. PNAS 2012 |
| I94L | 94 | 7.71 | 14.87 | 1.45 | 1.44 | Tamer et al. MBE 2019 |
| R98P | 98 | 2.82 | 34.93 | 0.41 | 4.65 | Tamer et al. MBE 2019 |
| L112V | 112 | 11.45 | 4.89 | | | Bershtein et al. PNAS 2012 |
| T113V | 113 | 32.90 | 21.40 | 0.50 | 1.10 | Fierke and Benkovic Biochemistry 1989 |
| I115V | 115 | 10.26 | 4.93 | | | Bershtein et al. PNAS 2012 |
| I115A | 115 | 7.59 | 15.18 | | | Bershtein et al. PNAS 2012 |
| G121V | 121 | 0.30 | 6.10 | 0.01 | 0.60 | Reynolds et al. Cell 2011 |
| W133F | 133 | 13.45 | 1.98 | | | Bershtein et al. PNAS 2012 |
| A145T | 145 | 10.00 | 2.78 | | | Bershtein et al. PNAS 2012 |
| F153S | 153 | 5.62 | 11.32 | 1.22 | 3.29 | Tamer et al. MBE 2019 |
| I155T | 155 | 11.27 | 4.14 | | | Bershtein et al. PNAS 2012 |
| I155L | 155 | 12.90 | 2.52 | | | Bershtein et al. PNAS 2012 |
| I155A | 155 | 11.95 | 4.41 | | | Bershtein et al. PNAS 2012 |
| WT | WT | 7.95 | 1.10 | 0.38 | 0.20 | Reynolds et al. Cell 2011 |

**Supplemental Table 1:** Steady-state DHFR kinetics parameters. Mean and standard deviation are reported across N=3 replicates when available.

| TYMS | kcat (s-1) | Km (µM) | kcat_std | Km_std |
|---|---|---|---|---|
| WT | 2.9 | 5.28 | 0.29 | 0.83 |
| R127A | 0.68 | 15.2 | 0.18 | 10.9 |
| Q33S | 1.99 | 4.43 | 0.19 | 1.8 |
| R166Q | 0.001* | 10* | 0 | 0 |

**Supplemental Table 2:** Steady-state TYMS kinetics parameters. Mean and standard deviation are reported across N=3 replicates when available. The signal-to-noise ratio for spectrophotometric measurements of R166Q TYMS activity is too low (given the slow rate of reaction) to allow accurate steady-state kinetics measurements. In this case, we assigned an arbitrarily slow $k_{cat}$ and a higher $K_m$. We observed that the model fit quality did not change appreciably for other values of R166Q provided they were substantially slower than WT.

| Dataset | # DHFR variants | # TYMS variants | # Fit Parameters | Ftot (µM) | [DHFR] (µM) | [WT TYMS] (µM) | [R127A TYMS] (µM) | [Q33S TYMS] (µM) | [R166Q TYMS] (µM) | K | n | g_max (per hour) | g_min (per hour) |
|---|---|---|---|---|---|---|---|---|---|---|---|---|---|
| Growth and metabolomics data | 5 | 2 | 8 | 8.54 +/- 3.7 | 0.32 +/- 0.04 | 0.12 +/- 0.01 | not in data | not in data | 77.29 +/- 7.76 | 0.04 | 1.58 | 0.7 | 0.48 |
| Plate-reader growth rates | 7 | 4 | 10 | 95.43 (+/- 19.06) | 3.16 +/- 3.91 | 0.08 +/- 0.10 | 0.23 +/- 0.27 | 0.28 +/- 0.34 | 21.18 +/- 5.47 | 0.65 +/- 0.25 | 0.62 +/- 0.35 | 1.41 +/- 0.21 | 0.2 +/- 0 |
| NGS-based growth rates | 38 | 3 | 9 | 47.61 (+/- 26/22) | 3.12 +/- 1.82 | 3.78 +/- 2.40 | not in data | 7.07 +/- 3.90 | 52.31 +/- 31.65 | 0.29 +/- 0.15 | 0.85 +/- 1.37 | 1.43 +/- 0.16 | 0.12 +/- 0.09 |

**Supplemental Table 3:** Model parameter fits for all three iterations of model fitting.

| Epistatic Cluster | TYMS | Positions |
| --- | --- | --- |
| negative | Q33S | 2+5+9+13+17+20+24+28+31 |
| negative | R166Q | 85 |
| positive | Q33S | 10+16+18+25+40+45+51+52+53+55+56+58+59+63+64+65+66+67+68+69+70+71+72+73+74+75+76+79+80+82+83+84+85+86+87+88+90+91+93+98+99+101+102+103+105+106+108+109+110+111 +112+115+117+120+136+139+141+143+145+146 |
| positive | R166Q | 2+3+5+8+9+10+11+12+13+16+17+18+20+21+23+24+25+26+28+29+32+33+38+39+42+44+45+47+48+49+50+51+52+53+55+56+58+59+60+61+62+63+64+65+66+67+69+70+71+72+73+74+75+76+ 79+80+82+88+90+91+92+94+97+98+99+101+102+104+108+109+111+112+115+117+119+120+126+128+131+132+147+149+150+153 |
| strong positive | Q33S | none |
| strong positive | R166Q | 4+6+7+14+15+31+35+40+41+46+54+81+93+100+103+110+121+122+123+125+133+148+151 |
| no significant epistasis | Q33S | 3+4+6+7+8+11+12+14+15+19+21+22+23+26+27+29+30+32+33+34+35+36+37+38+39+41+42+43+44+46+47+48+49+50+54+57+60+61+62+77+78+81+89+92+94+95+96+97+100+104+107+113+114 +116+118+119+121+122+123+124+125+126+127+128+129+130+131+132+133+134+135+137+138+140+142+144+147+148+149+150+151+152+153+154+155+156+157+158+159 |
| no significant epistasis | R166Q | 19+22+27+30+34+36+37+43+57+68+77+78+83+84+86+87+89+95+96+105+106+107+113+114+116+118+124+127+129+130+134+135+136+137+138+139+140+141+142+143+144+145+146+152+154 +155+156+157+158+159 |

**Supplemental Table 6**. Epistatic residue groups in DHFR as determined by k-means clustering. Plus signs are used as separators to facilitate creating selections in PyMOL.

| Conservation Cutoff | Number of conserved positions | Number of catalytic positions | p-value |
|---|---|---|---|
| 1.89 | 23 | 34 | 3.7 E -9 |
| 1.54 | 36 | 34 | 7.1 E -11 |
| 1.49 | 40 | 34 | 1.3 E -10 |
| 1.38 | 49 | 34 | 4.8 E -10 |

**Supplemental Table 7.** Statistical association between evolutionary conservation and DHFR positions where mutations systematically impact catalysis. All p-values were calculated by two-sided Fisher's exact test; we considered four cutoffs for evolutionary conservation (quantified by the Kullback-Leibler relative entropy) as previously reported[1]. DHFR positions associated to catalysis were defined by computing the average $\log_{10}$ effect on catalytic power across all mutations at a position; positions with an average mutational effect more than half a standard deviation below the mean were classified as linked to catalytic activity.
